# Distances and their visualization in studies of spatial-temporal genetic variation using single nucleotide polymorphisms (SNPs)

**DOI:** 10.1101/2023.03.22.533737

**Authors:** Arthur Georges, Luis Mijangos, Andrzej Kilian, Hardip Patel, Mark Aitkens, Bernd Gruber

**Affiliations:** Institute for Applied Ecology, University of Canberra, ACT 2601, Australia; Diversity Arrays Technology, University of Canberra, ACT 2601, Australia; National Centre for Indigenous Genomics, Australian National University, ACT 2601, Australia; RPS Australia Asia Pacific, Unit 2A, 45 Fitzroy Street, Carrington NSW 2294, Australia

**Keywords:** Principal Components Analysis, Principal Coordinates Analysis, Genetic distance, Metric distance, Euclidean distance, Structural variant

## Abstract

1. Distance measures are widely used for examining genetic structure in datasets that comprise many individuals scored for a very large number of attributes. Genotype datasets composed of single nucleotide polymorphisms (SNPs) typically contain bi-allelic scores for tens of thousands if not hundreds of thousands of loci.
2. We examine the application of distance measures to SNP genotypes and sequence tag presence-absences (SilicoDArT) and use real datasets and simulated data to illustrate pitfalls in the application of genetic distances and their visualization.
3. Euclidean Distance is the metric of choice in many distance studies. However, other measures may be preferable because of their underlying models of divergence, population demographic history and linkage disequilibrium, because it is desirable to down-weight joint absences, or because of other characteristics specific to the data or analyses. Distance measures for SNP genotype data that depend on the arbitrary choice of reference and alternate alleles (e.g. Bray-Curtis distance) should not be used. Careful consideration should be given to which state is scored zero when applying binary distance measures to sequence tag presence-absences (e.g. Jaccard distance).
4. Missing values that arise in the SNP discovery process can cause displacement of affected individuals from their natural groupings and artificial inflation of confidence envelopes, leading to potential misinterpretation. Filtering on missing values then imputing those that remain avoids distortion in visual representations. Failure of a distance measure to conform to metric and Euclidean properties is important but only likely to create unacceptable outcomes in extreme cases. Lack of randomness in the selection of individuals (e.g. inclusion of sibs) and lack of independence of both individuals and loci (e.g. polymorphic haploblocks), can lead to substantial and otherwise inexplicable distortions of the visual representations and again, potential misinterpretation.

## Introduction

Population genetics is the study of the interplay of genetic drift, gene flow, recombination and selection, and to a lesser extent mutation, as they come to influence the contemporary genetic composition of populations. Many measures of genetic similarity and dissimilarity have been developed to quantify inter-individual and inter-population variation arising from these processes. The concept of genetic distance (Sanghvi, 1953) is now a fundamental tool in genetics (Nei & Kumar, 2000).

Application of genetic distances in studies of spatio-temporal variation is common because substructuring of populations can arise from interesting combinations of genetic, demographic and biogeographic processes. One area that has received substantial attention is isolation by geographic distance (Wright, 1943) which, in a homogenous and continuous landscape, is perhaps the simplest instance of departure from panmixia in a widespread population. Isolation by distance is typically examined by comparing a genetic distance matrix with geographic distance between individuals, sub-populations or sampling localities. Fst and Chord Distance are often chosen to examine any relationship with geographic distance (Séré et al., 2017). Such a relationship is often used as the null hypothesis against which to examine more complicated hypotheses to explain patterns of variation across the landscape (Manel et al., 2003; Spear et al., 2015). In particular, isolation by distance can be complicated by variation in resistance to gene flow in complex landscapes (McRae, 2006). Introduction of the concept of landscape resistance delivers a continuous set of circumstances between genetic structure arising from isolation by distance in otherwise freely breeding populations and subdivision between populations of a species where gene flow is absent, rare or an episodic event. More novel applications of genetic distances include stratified sampling of a core collection of specimens from a larger pool of accessions to maximally capture genetic diversity (Rogers D: Jansen & van Hintum, 2007) and tracking the relationships of SARS-CoV-2 variants (Jaccard D: Yin, 2020). Another area of endeavour that has benefitted from the advances afforded distance measures is phylogeography, both in the context of human populations (Pugach & Stoneking, 2015; Tassi et al., 2015) and animal and plant populations (Edwards et al., 2015). Here, genetic substructuring is mapped onto the landscape and interpreted in the context of historical (Georges et al., 2018) and contemporary (Mijangos et al., 2022) barriers to geneflow. SNPs and distances derived from them are also useful in population assignment (Negrini et al., 2009) and for putatively identifying migrants within source populations (Mamoozadeh et al., 2020). These analyses rely on genetic distances and their visualization.

Genetic distances fall into several broad classes. Some are obtained by direct measurement, such as immunological distance (Faith, 1985) or DNA-DNA hybridization (de Ley et al., 1970; Hirayama et al., 1996; Kirsch et al., 1990). However, most genetic distances are calculated from character states (genotypes) arranged as a matrix of individuals (entities) by genetic loci (attributes) Genetic distances can be further classified by whether they will be used to infer patterns of ancestry and descent among species (phylogenetics), the structure and relationships among populations of a species at various scales of divergence with or without gene flow (population genetics), or relationships among individuals (e.g. kinship).

In this paper, we deal with distances defined for both individuals and populations and their visualization in the broader context of divergence among populations and subpopulations. The array of available measures of distance and similarity/dissimilarity is daunting (Deza & Deza, 2009). We specifically focus our attention to analyses of single nucleotide polymorphisms (SNPs), markers that are commonly used in studies of spatial or temporal variation among individuals and populations of a species or closely related species. We also consider genetic distances as they apply to complementary dominant sequence tag presence-absence markers (SilicoDArT markers, analogous to microarray DArTs first described by Jaccoud et al. (2001) but extracted in silico to sequences obtained from genomic representations e.g. Ali et al., 2020; Mahboubi et al., 2020; Elshibli and Korpelainen, 2021; Nantongo et al., 2022). Although genetic distances are used in a wide range of contexts (Jansen & van Hintum, 2007; Libiger et al., 2009; Yin, 2020), we also restrict our attention (not exclusively) to the application of distances in studies of spatial and temporal genetic structure among populations where allelic profiles are governed principally by recent or contemporary processes of drift, selection and gene flow. We do not consider distance measures used to reveal deeper historical patterns of ancestry and descent among lineages on independent evolutionary trajectories where the pattern of mutational events dominates (species-level phylogenetics). Nor do we consider distance measures used to quantify distances between SNPs loci themselves as opposed distances between individuals or populations (Müller et al., 2005).

Our treatment of suitable genetic distance measures is predicated on the criterion that they should agree with our commonsense notion of a distance. They should satisfy, to an adequate level of approximation, the properties of a metric distance (Legendre & Legendre, 2012). This can be particularly important when the genetic distances are to be represented graphically in a space defined conventionally by Cartesian coordinates and, in particular, Principal Coordinates Analysis (PCoA) (Gower, 1966). We also discuss Principal Components Analysis (PCA) (Pearson, 1901; Hotelling, 1933; Jolliffe, 2002) which, although it does not take a distance matrix as input, is a closely related technique that provides a foundational underpinning to PCoA. The two analyses are collectively referred to as ordination. These techniques have found application in many diverse fields such as ecology, economics, psychology, meteorology, oceanography, human health and genetics as a descriptive and an exploratory tool to generate hypotheses for further examination (Jolliffe & Cadima, 2016) rather than a formal statistical analysis (but see Patterson et al., 2006).

We examine some practical aspects of applying distance analyses and their visualization. We pay particular attention to underlying assumptions of the application of distance analyses and visualization of spatio-temporal structure using ordination, many of which are poorly appreciated. These assumptions include the properties of various distance measures and the importance of metric and/or Euclidean properties for visualization, the impact of missing values and the importance of randomness in sampling and independence of individuals (relatedness), their genotypes and the loci selected for screening.

Finally, we appreciate that PCA has recently come under criticism (Elhaik, 2022), but argue that many of these criticisms derive from the over-interpretation of the patterns of genetic variation revealed by PCA. We argue strongly that PCA and PCoA are exploratory tools that draw out patterns from multivariable data by aggregating correlated influences into a relatively few dimensions. These techniques have little to contribute to understanding of the underlying events and processes that led to those patterns. The patterns thus are suggestive but not demonstrative of underlying causes. Interpretations of the causes of the patterns need to be pursued by more definitive subsequent analysis, beyond the scope of this review, to distinguish between competing explanations for the observed patterns.

### Box 1. Glossary of terms

**Population**

A group of individuals of the same biological species that, by virtue of occupying a well-defined geographic provenance, have opportunity to interbreed. Changes in the allelic profiles of a subset of individuals will come to affect the whole.

**Sub-population**

A subset of individuals of a population, occupying a well-defined geographic provenance and which, because of resistance to geneflow arising from partial or episodic isolation, have a distinct allele frequency profile. For ease of communication, we refer to populations on the implicit understanding that the point also applies also to subpopulations.

**SNP**

Single Nucleotide Polymorphism. A single base pair mutation that is polymorphic.

**SNP genotype data**

SNPs scored across multiple loci for each individual. Codominant markers.

**SilicoDArT data**

Presence or Absence of the amplified sequence tags across multiple loci for each individual. Dominant markers scored as 0 for absence and 1 for presence. Absences are typically null alleles. Hemizygous state scored as present (1).

**Entity**

Something with distinct and independent existence. May refer to a sample, subject, individual or population from which the genotypes are scored.

**Attribute**

A character that is scored for each entity, in this case the loci.

**State**

Refers to the value of an attribute (locus) for a given entity (individual) in the data matrix. Typically scored as 0, 1, 2 for SNP genotype data; 0, 1 for SilicoDArT data.

**Call Rate**

Measures the rate at which a SNP or SilicoDArT markers are successfully called (scored). They may not be called because of a mutation at one or both of the restriction enzyme recognition site or they may be missed in the sequencing because of insufficient read depth. Either way, the score is missing.

**Distance**

Term used loosely to encompass distance and dissimilarity measures (Gower & Legendre, 1986).

**Euclidean distance**

Metric distance that further can be represented faithfully in a space defined by Cartesian coordinates.

**Metric distance**

Distance that satisfies the formal metric properties – positive definite, symmetric, and conforms to the triangle inequality.

**Dissimilarity measure**

A quantitative measure of how different two individuals or populations are. Does not necessarily conform to the properties of a metric.

**PCA**

Principal Components Analysis, a classical ordination technique drawing upon the covariance matrix (unstandardized) or the correlation matrix (standardized) for the entities by attributes matrix.

**PCoA**

Principal Coordinates Analysis, a generalization of PCA that admits a wider range of distance and dissimilarity measures.

**Reference allele**

The allele, typically the most frequent or major allele, that is arbitrarily chosen as reference to distinguish it from the alternate allele. Scored as 0 in the homozygous state.

**Alternate allele**

The allele, typically the least frequent or minor allele, that is arbitrarily chosen as alternate to distinguish it from the reference allele. Scored as 2 in the homozygous state.

**MDS**

Multidimensional scaling whereby individuals are represented in a Cartesian space of a specified dimension by iteratively matching the distances in that space with those of the associated distance or dissimilarity measure. It can be applied to data converted to ranks.

## Datasets and Methods

A dataset was constructed from a SNP matrix generated for the freshwater turtles in the genus *Emydura* (Georges et al., 2018), a recent radiation of Chelidae in Australasia. The dataset includes selected populations that vary in level of divergence to encompass variation within species and variation between closely related species. Sampling localities with evidence of admixture between species were removed. Monomorphic loci were removed, and the data was filtered on call rate (>95%), repeatability (>99.5%) and read depth (5*x* < read depth < 50*x*). Where there was more than one SNP per sequence tag, only one was retained at random. The resultant dataset had 18,196 SNP loci scored for 381 individuals from 7 sampling localities or populations – *Emydura victoriae* [Ord River, NT, n=15], *E. tanybaraga* [Holroyd River, Qld, n=10], *E. subglobosa worrelli* [Daly River, NT, n=25], *E. subglobosa subglobosa* [Fly River, PNG, n=55], *E. macquarii macquarii* [Murray Darling Basin north, NSW/Qld, n=152], *E. macquarii krefftii* [Fitzroy River, Qld, n=39] and *E. macquarii emmotti* [Cooper Creek, Qld, n=85]. The missing data rate was 1.7%, subsequently imputed by nearest neighbour (Beretta & Santaniello, 2016) to yield a fully populated data matrix. The data are a subset of those published by Georges et al. (2018) for illustrative purposes only. A companion SilicoDArT dataset for the same individuals was also generated, filtered on read depth (*>* 12x), repeatability (>99.5%), and call rate (>95%).

The above manipulations were performed in R package dartR (Gruber et al., 2018; Mijangos et al., 2022; refer supplementary text for the R Code). Principal Components Analysis was undertaken using the glPCA function of the R adegenet package (as implemented in dartR, Gruber et al., 2018; Jombart, 2008; Jombart & Ahmed, 2011). Principal Coordinates Analysis was undertaken using the pcoa function in R package ape (Paradis & Schliep, 2019) implemented in dartR (Mijangos et al., 2022).

To exemplify the effect of missing values on SNP visualisation using PCA (Figure 7), we performed computer simulations in dartR (Mijangos et al., 2022). Briefly, we simulated ten populations that reproduced over 200 non-overlapping generations. Simulated populations were placed in a linear series with low dispersal between adjacent populations (one disperser every ten generations). Each population had 100 individuals, of which 50 individuals were sampled at random. Genotypes were generated for 1,000 neutral loci on one chromosome. We then randomly selected 50% of genotypes and set them as missing data. Principal Components Analysis was undertaken using the glPCA function of the R adegenet package (as implemented in dartR, Gruber et al., 2018; Jombart, 2008; Jombart & Ahmed, 2011). R script and input files to replicate this analysis are provided in the supplementary text.

The data for the Australian Blue Mountains skink *Eulamprus leuraensis* (Figure 8) were generated for 372 individuals collected from 17 swamps isolated to varying degrees in the Blue Mountains region of New South Wales. Tail snips were collected and stored in 95% ethanol. The tissue samples were digested with proteinase K overnight and DNA was extracted using a NucleoMag 96 Tissue Kit (MachereyNagel, Duren, Germany) coupled with NucleoMag SEP (Ref. 744900) to allow automated separation of high-quality DNA on a Freedom Evo robotic liquid handler (TECAN, Miinnedorf, Switzerland). SNP data were generated by the commercial service of Diversity Arrays Technology Pty Ltd (Canberra, Australia) using published protocols (Georges et al., 2018; Kilian et al., 2012; Sansaloni et al., 2011). A total of 13,496 loci were scored which reduced to 7,935 after filtering out secondary SNPs on the same sequence tag, filtering on reproducibility (threshold 0.99) and call rate (threshold 0.95), and removal of monomorphic loci. The resultant data is used to demonstrate the impact of a substantial inversion on the outcomes of a PCA.

To test the effect of having closely related individuals (parents and offspring) on the PCoA pattern we ran a simulation using dartR (Gruber et al. 2018, Mijangos et al. 2022), where we picked up two individuals to become the parents with 2-8 offspring. We ran a PCoA for all of the simulated cases (Figure 9 shows the case of 0 and 8 offspring). R script and input files to replicate this analysis are provided in the supplementary text.

### The Concept of Distance

Standard Euclidean distance is a common-sense notion derived as an abstraction of physical distance. Applying Standard Euclidean distance to SNP data is straightforward. The horizontal axis of Figure 1a represents SNP Locus 1 and the values *x*_*A1*_ and *x*_*B1*_ represent the scores for that locus (0 or 1 or 2 for SNPs; 0 or 1 for tag presence-absence) for individuals A and B respectively; the vertical axis represents SNP Locus 2 with the values *x*_*A2*_ and *x*_*B2*_ similarly defined.

**Figure 1.**
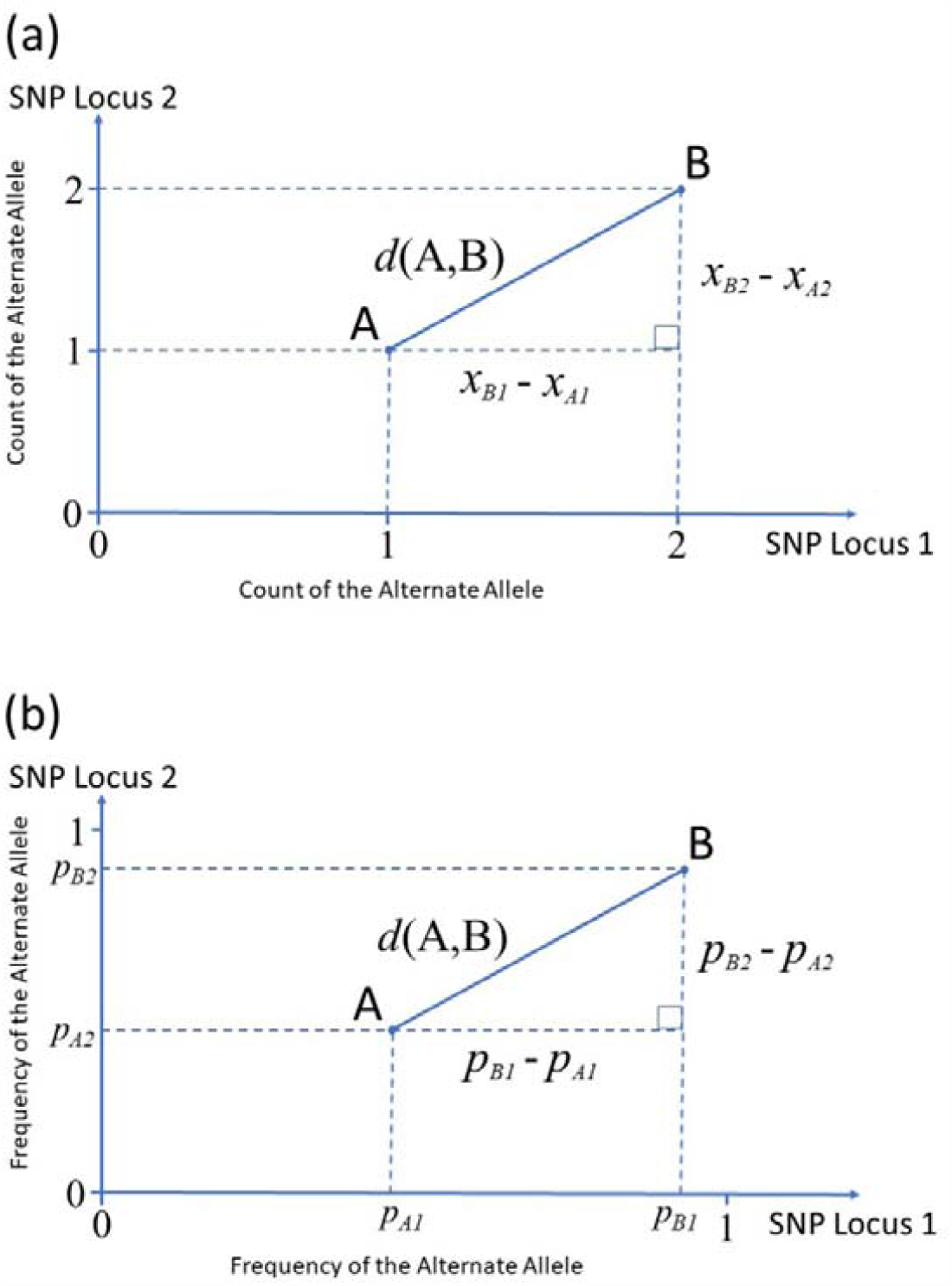
Distance between two points A and B represented in two-dimensional space can be calculated from their Cartesian coordinates using Pythagoras’ rule – the square of the hypotenuse of a right-angled triangle is equal to the sum of the squares of the two adjacent sides. Each axis can be considered to represent a locus (a), with the value taken by an individual (A or B) on that axis called as *x*_*Ai*_ and *x*_*Bi*_ = 0, 1 or 2 for SNP genotype. As such, each individual is represented by a point in a multidimensional space defined by the *i* = 1 to *L* loci. Populations A and B can be similarly depicted, as in (b), with the relative frequency of the alternate allele in the population (*p*_*Ai*_ and *p*_*Bi*_) as the value taken on each SNP axis. Representation for SNP PA data can be similarly defined.

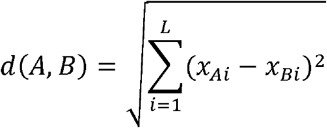

Standard Euclidean Distance can be similarly defined for populations,

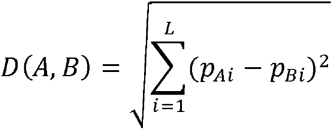

where *p*_*Ai*_ and *p*_*Bi*_ are the relative frequencies of the alternate allele at locus *i* in populations A and B respectively (Figure 1b).

The equations for *d(A,B)* and *D(A,B)* can be scaled to fall in the range [0,1] on noting that its maximum is achieved for SNP genotype data when all *x*_*Ai*_ are 0 and all *x*_*Bi*_ are 2; for SilicoDArT data the maximum is achieved when all *x*_*Ai*_ are 0 and all *x*_*Bi*_ are 1. Note also that the equations for *d(A,B)* and *D(A,B)* are symmetric with respect to choice of which allelic state is assigned to reference (0 for homozygous reference) and which is assigned to alternate (2 for homozygous alternate). Interchanging the reference and alternate alleles (that is applying transformation *x’ = 2 – x* for SNP genotype data or *x’ = 1 – x* for SilicoDArT data) has no impact on the value of the distance between A and B.

A Mantel Test (Guillot & Rousset, 2013; Mantel, 1967) can be used to compare two genetic distance matrices (or a genetic distance and a geographic distance matrix) provided they are constructed for the same entities and those entities are independent (i.e. not autocorrelated).

### Generalization of the concept of Distance

Standard Euclidean distance is just one of many distance measures. The concept of distance more generally can be distilled down to three basic properties. For a metric distance we have:

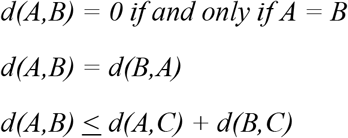

The first condition asserts that indiscernible entities are one and the same. The second condition asserts symmetry. The last condition is referred to as the triangle inequality which enforces the notion that the distance between two points is the shortest path between them. From these properties we can conclude that metric distances must be non-negative.

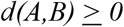

In essence, metric distances are well-behaved distances. Standard Euclidean Distance, as with all Euclidean distances, is a metric distance.

Graphically, the metric properties make complete sense for a distance (Figure 2). Given three points defined by the distances between them, the position of each of them is defined (Figure 2a). This is necessary (though not sufficient) if we are to represent our distances in Cartesian coordinate space a fundamental requirement of PCoA.

**Figure 2.**
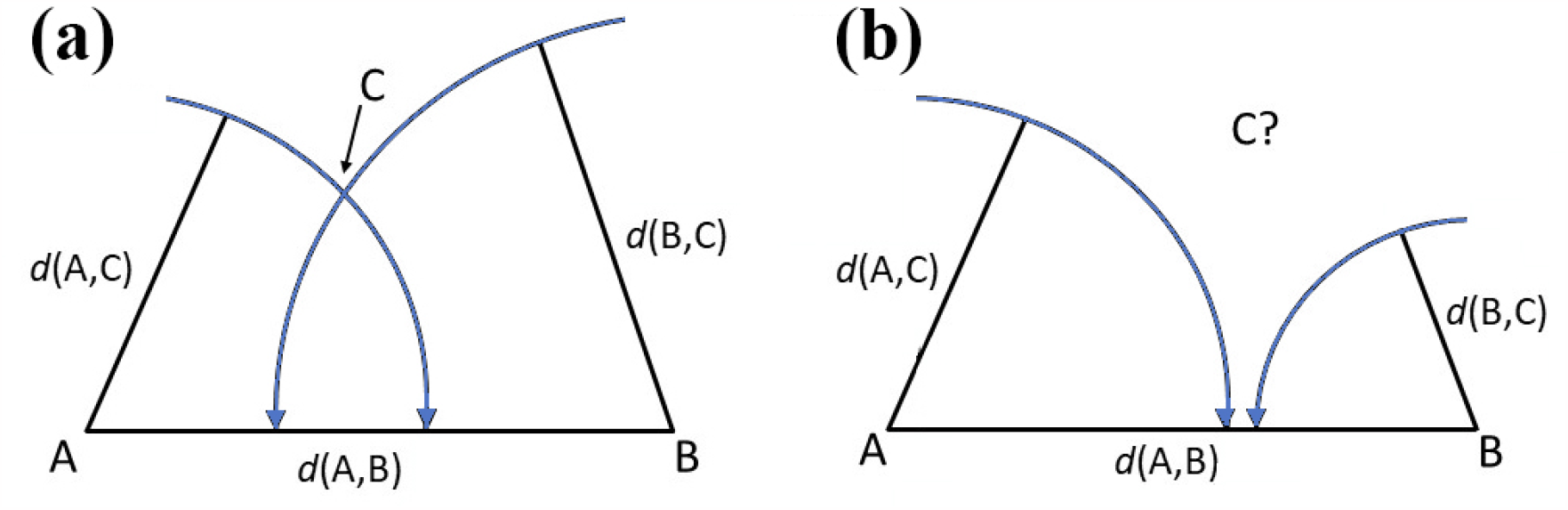
Visual representation of the triangle inequality as used to define a metric distance. **(a)** – if three distances between points A, B and C satisfy the metric property *d(A, B) d(A, C)* + *d(B, C)*, then the position of each of A, B and C in space is well defined – when the lines represented by d(A,C) and d(B,C) are swung through an arc (blue lines), they meet to define a unique point C **(b)** – if not, and *d(A, B) > d(A, C)* + *d(B, C)*, then point C is undefined – when the lines represented by d(A,C) and d(B,C) are swung through an arc, they fail to meet and so C is undefined by the information contained in the distances.

While the metric properties of a distance are clearly important, many measures used in genetics are non-metric. An example of a dissimilarity measure that fails to satisfy the symmetry condition is one defined on private alleles. Private alleles are those uniquely possessed by one population when compared to other populations. The number of private alleles possessed by population A compared to population B will typically be different from the number of private alleles possessed by population B compared with A. Thus, the resultant distances will not satisfy the second metric criterion of symmetry. Other genetic distances in use do not satisfy the triangle inequality. For example, calculating the percentage of fixed allelic differences between two entities (the percentage of alleles that are homozygous for the reference allele in one entity and homozygous for the alternate allele in the other entity) as a measure of distance satisfies the first two conditions of a metric distance, but not the triangle inequality and so is non-metric. Nor is Nei’s D (Nei, 1972) a metric distance for the same reason (Farris, 1981), but the common alternative of Rogers’ D (Rogers, 1972) is metric (essentially equivalent to Euclidean Distance). *F*_*ST*_ is non-metric (Abisser and Rosenberg, 2020). The Bray-Curtis dissimilarity measure is non-metric but is rank-order similar to the Jaccard distance, which is metric. And so on (refer Legendre & Legendre, 2012--tables 7.2 and 7.3).

**Table 1.** Some genetic distances commonly applied to binary data and derived from sequence tag presence or absence, such as to SilicoDArT data. Variables are described in the text. Formulae illustrate the adjustments made to the denominator for Jaccard and Sørensen distances rather than being presented in their conventional algebraic form. Software packages in R for calculating distances for binary data, including those listed above using the same terminology, include vegan (Oksanen et al., 2014) and dartR (Gruber et al., 2018; Mijangos et al., 2022).

| Name | Formula | Alternate Names | Reference |
| --- | --- | --- | --- |
| Standard Euclidean Distance | $d_E = \sqrt{\frac{N_{01} + N_{10}}{L}}$ | | |
| Simple Matching Distance | $d_{SM} = \frac{N_{01} + N_{10}}{L}$ | Scaled Hamming Distance | (Sokal & Michener, 1958) |
| Jaccard Distance | $d_J = \frac{(N_{01} + N_{10})}{L - N_{00}}$ | Marczewski-Steinhaus D (Marczewski & Steinhaus, 1959); Ružička D; Soergel D | (Jaccard, 1912) |
| Sørensen Distance | $d_s = \frac{(N_{01} + N_{10})}{L - N_{00} + N_{11}}$ | Dice D (Dice, 1945) | Sørensen D (Sørensen, 1948) |

**Table 2.**
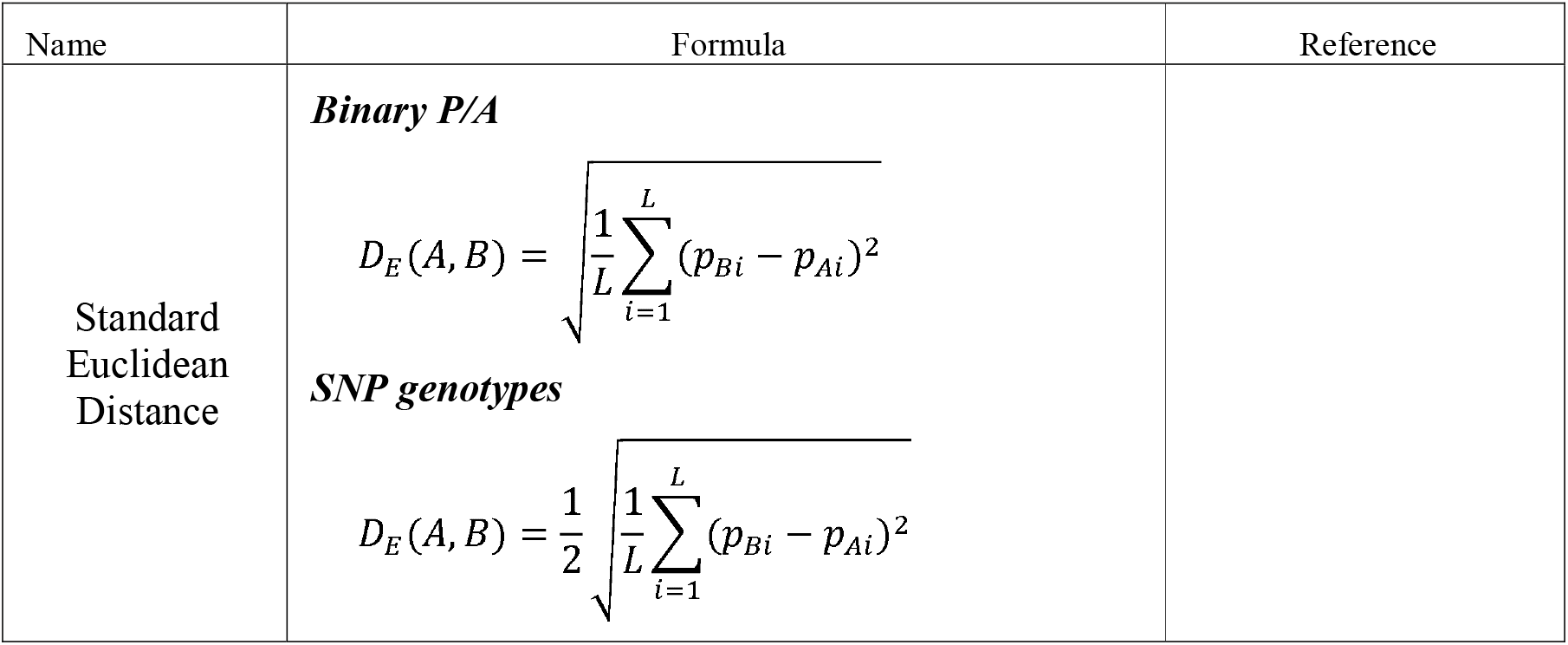
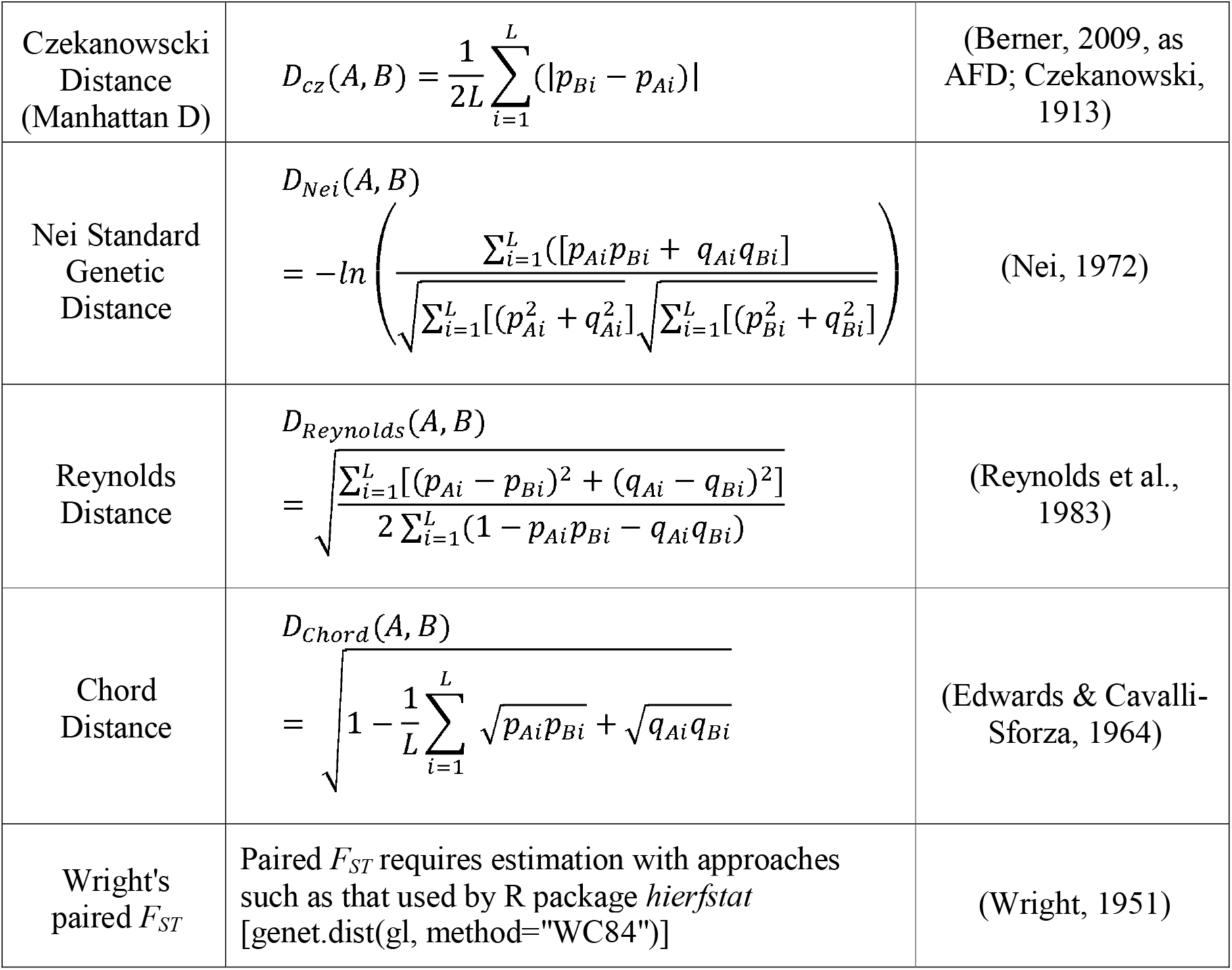
Some genetic distances commonly applied to populations. *p*_*Ai*_ is the proportion of the alternate allele for Locus *i* in population A, *p*_*Bi*_ is the proportion of the alternate allele for Locus *i* in population B. *q*_*Ai*_ and *q*_*Bi*_ are similarly defined for the reference alleles. *L* is the number of called loci. Software packages in R for calculating distances for binary data, including those listed above using the same terminology, include vegan (Oksanen et al., 2014), adegenet (Jombart, 2008), poppr (Kamvar et al., 2014), pegas (Paradis, 2010) and dartR (Gruber et al., 2018; Mijangos et al., 2022). Package heirfstat (Goudet, 2005) is the software of choice for calculating F statistics.

### Genetic Distances for Individuals

#### Binary Data

SilicoDArT (sequence tag presence-absence) data involves scoring loci as “called” [1] or “not called” [0]. They are called because if the two restriction enzymes (used in the accompanying DArTSeq or ddRAD) find their mark, the corresponding sequence tags are amplified and sequenced, and the locus is scored as 1 for that locus. If, however, there is a mutation at one or both of the restriction enzyme sites, then the restriction enzyme(s) does not find its mark, the corresponding sequence tag in that individual is not amplified or sequenced (null allele), or if it is amplified from a different start site, is no longer considered homologous during pre-processing. The individual is scored as 0 for that locus. SilicoDArT markers are commonly used in an agricultural setting, to generate datasets that are complementary to SNP datasets (Mahboubi et al., 2020; Nantongo et al., 2022). They have less commonly been used in the context of population genetics (but see Elshibli and Korpelainen, 2021).

Unlike the co-dominant SNP genotype markers, SilicoDArT markers are dominant markers. Technical artefacts such as low DNA quality or quantity, or failure of experimental protocol resulting in shallow sequencing depth, may occasionally lead to missing data which will be erroneously interpreted as a null allele. Typically, this is overcome by quality control processes. Specific preprocessing options are used to separate true null alleles (sequence tag absences) from sequence tags present in the target genome but missed by the sequencing because of low read depth. First, the sequence tags are filtered more stringently on read depth than for SNPs, typically with a threshold of 8*x* or more. Second, the distributions of the sequence tag counts are examined for bimodality (clustering on either 0 or 1) and those loci showing a continuum of counts rather than two clusters are discarded. Hemizygous loci are scored as missing (-). Finally, the scores are tested for reproducibility using technical replicates, providing the analyst with the opportunity to filter loci with poor reproducibility in the SilicoDArT scores. In this way, a reliable set of dominant markers are obtained, to complement the SNP dataset generated from the same individuals. Note that although the SilicoDArT and SNP genotypes are derived from the same samples and representation, the SilicoDArT markers include sequence tags that do not have a SNP, that is, sequence tags that are invariant across all individuals.

Many binary distance measures (Deza & Deza, 2009) have been co-opted for genetic studies using these dominant markers (e.g. Jaccard Distance, Ali et al., 2020). The calculation of genetic distance from SilicoDArT data involves four intermediate sums. Taking two individuals A and B, we can count the different cases,

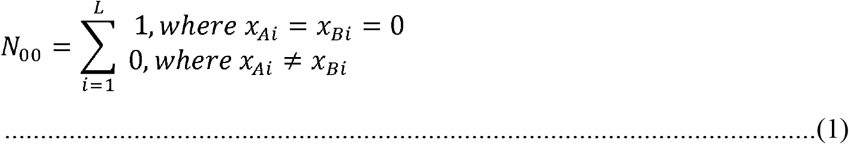

where *x*_*Ai*_ and *x*_*Bi*_ are the SilicoDArT scores for individuals A and B respectively. *N*_*00*_ is the sum of loci scored 0 (absent) for both individuals;

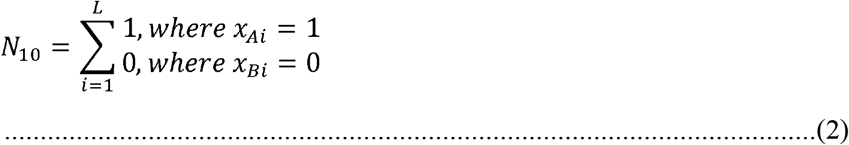

that is, sum loci scored 1 (present) for Individual A and 0 (absent) for Individual B;

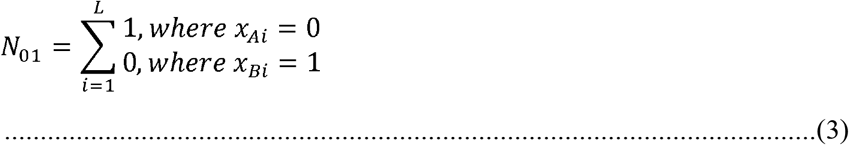

that is, sum of loci scored 0 (absent) for Individual A and 1 (present) for Individual B;

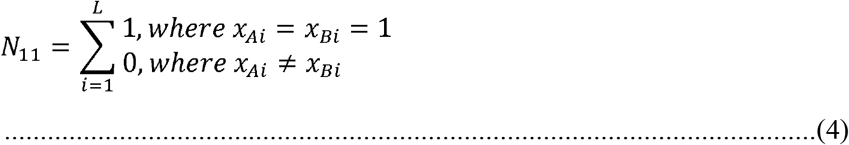

that is, sum of loci scored 1 (present) for both individuals. These summations do not include loci for which data are missing (NA) for one or both individuals.

The number of loci L is given by

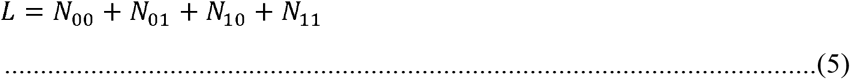

Based on the above intermediates [(1) – (5)], there are several ways to calculate a binary distance between two individuals (Choi et al., 2010), some of which are shown in Table 1. The two most commonly used distance measures are Standard Euclidean Distance and Simple Matching Distance (Sokal & Michener, 1958). Both are based on the number of mismatches between two individuals (*N*_*01*_ *+ N*_*10*_), expressed as a proportion of the total number of loci considered (*L*), and used when there is symmetry (equivalence) in the information carried by 0 (absence) and 1 (presence). Simple Matching Distance is simply the Standard Euclidean distance squared and, as such, will give greater weight to larger distances than Euclidean distance. This might be of value when looking for clusters of entities in subsequent analyses. Simple Matching Distance is a metric distance (Orlóci, 1978:62).

Jaccard Distance is a variation on the Simple Matching Distance that down-weights joint absences and so is no longer symmetric with respect to 0 and 1 scores. Note that multiple independent mutations can result in a 0 score for a sequence tag, and absences of a sequence tag (again 0) may arise from a failure to detect; both can contribute artificially to the similarity between two entities. One might therefore consider recoding the data (*x’ = 1 -x*) so that 1 represents presence of a mutation at one or both of the restriction enzyme sites (i.e. absence of the amplified tag) and 0 represents absence of such a disruptive mutation (i.e. success in amplifying the sequence tag). Having made this simple adjustment, the Jaccard Distance will down-weight joint absence of a disruptive mutation. The Jaccard Distance is a metric distance (Levandowsky & Winter, 1971).Sørensen Distance adjusts the denominator to down-weight the joint absences (0,0) and up-weight joint presences (1,1) (Table 1) and so is a stronger response to the uncertainty surrounding joint absences that in the Jaccard Distance. As with the Jaccard Distance, you might consider reversing the scores for absence (0) and presence (1) to 1 and 0 respectively when dealing with SilicoDArT data. Special attention may be required to manage missing values when applying the Sørensen and Jaccard Distances, because adjustment of the denominator in their formulae (Table 1) can lead to a potential systematic bias (Orlóci, 1978:62). The Sørensen Distance is not a metric distance (Orlóci, 1978:61).

#### SNP Genotype Data

SNP data take on three values at a locus: 0 for homozygous reference allele; 1for heterozygous; and 2 for homozygous alternate allele, that is, the count of the alternate allele. Software packages PLINK (Chang et al., 2015), SNPTEST (Marchini et al., 2007) and dartR (Gruber et al., 2018; Mijangos et al., 2022) use this scoring scheme as the default. The alternative scoring scheme of 0 for homozygous reference allele, 1 for homozygous alternate allele and 2 for heterozygotes is commonly used in genome-wide association studies (GWAS), and software such as GCTA (Genome-wide Complex Trait Analysis, Yang et al., 2011) and by Diversity Arrays Technology Pty Ltd (Canberra, Australia).

SNPs have particular characteristics that influence the choice of a distance measure. SNPs can have more than two alleles (i.e. are multiallelic) but in practice, sites with more than two allelic states are filtered out during the selection of markers as part of pipelines to eliminate non-homologous sequence tags. Fortunately, multiallelic SNP sites are rarely observed. As a result of this filtering, SNP markers are bi-allelic and have characteristics that substantially limit options for a distance measure. For example, the values of 0 and 2 carry equal weight because the choice of reference allele and alternate allele is arbitrary. Any distance measure that gives differential weight to joint zeros (e.g. Bray & Curtis, 1957 Distance) will yield values that depend on the arbitrary choice of which allele is reference and which is alternate at each locus. Such distance measures can be eliminated from options available for SNP genotype data. Standardization or normalization across attributes (loci) is not required because the attributes (loci) are all measured as genotypes on the same scale (0, 1 or 2); this is not the case under some circumstances in multiallelic systems. That said, the biallelic nature of SNP markers permits the easy calculation of the maximum distance possible, which permits scaling distances to fall conveniently in the range of [0,1] while maintaining comparability across studies. Finally, issues that arise in a multiallelic context do not apply in a biallelic context. For example, Rogers D (Rogers, 1972), and therefore Standard Euclidean Distance, can yield undesirable results when two populations are both polymorphic at a site, but share no alleles (Nei & Kumar, 2000:246). This situation does not arise in the biallelic case, and so Standard Euclidean Distance (or equivalently Rogers D) is often the distance of choice in SNP studies for both individuals and populations.

Standard Euclidean Distance as applied to SNP genotype data is defined in the usual way with loci as the axes in a coordinate space, and the value on each axis is 0, 1 or 2 as defined above. The scaling factor of ½ arises because the maximum squared distance between two individuals at a locus is (2-0)^2^ = 4.

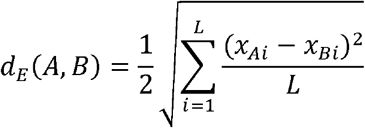

where *x*_*Ai*_ and *x*_*Bi*_ are the counts of the alternate allele at locus *i* for individual A and B respectively; *L* is the number of loci for which both *x*_*Ai*_ and *x*_*Bi*_ are non-missing.

The Simple Matching Distance uses the counts of shared alleles between two individuals *i* and *j* at a locus is given by

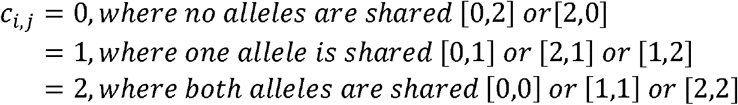

and is calculated as

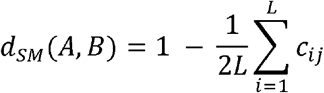

where *L* is the number of loci non-missing for both individuals *i* and *j*. It is metric and similar to the Allele Sharing Distance (Gao & Starmer, 2007), differing from it by a factor of 2. Simple Matching Distance lacks a square and so does not give greater weight to larger distances as does Standard Euclidean Distance.

Czekanowski Distance (Czekanowski, 1913) is calculated by summing the scores across the axes

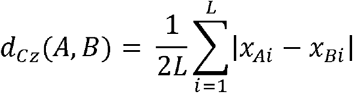

where *x*_*Ai*_ and *x*_*Bi*_ are the counts of the alternate allele at locus *i* for individual A and B respectively; *L* is again the number of loci for which both *x*_*Ai*_ and *x*_*Bi*_ are non-missing. Referred to also as the Manhattan D or the City Block D, Czekanowski Distance is a metric distance. Czekanowski Distance is often preferred over Standard Euclidean distance when there are extreme outliers among the data.

Software packages in R for calculating distances for SNP genotypes, including those listed above using the same terminology, include vegan (Oksanen et al., 2014), adegenet (Jombart, 2008), poppr (Kamvar et al., 2014), pegas (Paradis, 2010) and dartR (Gruber et al., 2018; Mijangos et al., 2022).

### Genetic Distances for Populations

Standard Euclidean Distance as applied to allele frequencies is defined in the usual way with loci (attributes) represented by the axes in a coordinate space, and the values (states) for each population (entity) on each axis set to the frequency of the alternate allele for the respective locus (Table 2). An appropriate scaling factor is applied to bring the value of the distance into the range [0,1], which renders it identical to Roger’s D (Rogers, 1972). Standard Euclidean distance is a model-free metric distance, in that its formulation makes no assumptions regarding evolutionary processes generating the genetic distances.

Nei’s standard genetic distance (Nei, 1972) is favoured by some over Standard Euclidean Distance because of its relationship to divergence time. When populations are in mutation-drift balance throughout the evolutionary process and all mutations result in new alleles in accordance with the infinite-allele model, Nei’s *D* is expected to increase in proportion to the time after divergence between two populations (Nei, 1972). Nei’s *D* is non-metric.

Reynolds genetic distance (Reynolds et al., 1983) is also approximately linearly related to divergence time in theory, but unlike Nei’s Standard Genetic Distance, it is based solely on a drift model and does not incorporate mutation. Thus, it may be more appropriate than Nei’s distance for spatial population genetics (divergence on relatively short timescales) and in particular, representation of genetic similarity in trees (phenograms) or networks where branch lengths need to be interpretable. A better approximation (Reynolds et al., 1983) of the linear relationship with time is given by

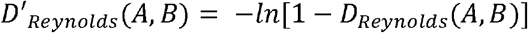

Chord Distance (Edwards & Cavalli-Sforza, 1964) (Table 2) assumes divergence between populations is via drift alone, and so again may be more appropriate than Nei’s *D* for spatial population genetics. Chord Distance is based on Geodesic or Angular Distance (Bhattacharyya, 1946; Edwards, 1971; Edwards & Cavalli-Sforza, 1967),

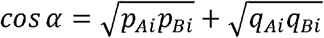

where α is the Angular Distance. For a geometric interpretation using biallelic data, see Nei & Kumar (2000:267). Chord Distance approximates Angular Distance by replacing the arc distance with the length of the corresponding straight-line segment (Edwards & Cavalli-Sforza, 1964) (Table 2). It is a metric distance that can be transformed to be approximately Euclidean (Edwards, 1971). As with Reynold’s *D*, Chord Distance is proportional to shallow divergence time (for relatively small values of D) under specified assumptions (Edwards, 1971). It may be preferred over the model-free Standard Euclidean Distance where an underlying genetic model of divergence with time is preferred.

Bray-Curtis Distance applies to abundances and down-weights joint absences in a manner analogous to the Jaccard Distance. There is some confusion arising from the translation of Bray-Curtis Distance (Bray & Curtis, 1957) from an ecological perspective to a genetics perspective. When applied to SNP data, the Bray-Curtis Distance uses the abundance of the alternate allele which renders it asymmetric with respect to the arbitrary choice of which is reference and which is alternate allele. Bray-Curtis Distance defined in this way should thus not be used on SNP data. Some authors have suggested that the Bray-Curtis equation be applied separately to the counts of each allele, averaged for the locus, for computing genetic distance between individuals or populations (Allele Frequency Difference or AFD, Berner, 2009; Bray-Curtis/Allele Frequency Difference or BCAFD, Sherwin, 2022). However, when this formulation is applied to either individuals or populations, it becomes algebraically equivalent to the Czekanowski Distance (=Manhattan D). It no longer has the properties of Bray-Curtis Distance (having been rendered metric and symmetric). To avoid confusion, members of this class of distances should not be referred to as Bray-Curtis, AFD or BCAFD, but rather as Czekanowski Distance or the more familiar Manhattan D, given that naming of a distance measure should not be context dependent. The same issue has arisen in ecology (Yoshioka, 2008). Czekanowski Distance applied to SNP data and corrected for maximum value dependency, referred to as ^A^A Distance (Sherwin, 2022), restores the asymmetry with respect to choice of reference and alternate allele.

Wright’s *F-statistics* (Wright, 1951) describe the distribution of genetic diversity within and between populations (Holsinger & Weir, 2009). *F-statistics* are defined in terms of the proportion of heterozygotes observed (*H*_*obs*_) and the proportion of heterozygotes expected under Hardy-Weinberg equilibrium (*H*_*exp*_), as follows:

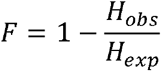

A deficit in the observed proportion of heterozygotes compared with that expected under Hardy-Weinberg equilibrium will yield a Wright’s *F* that deviates from zero. If, at a single locus, observed and expected heterozygosity are in agreement, then *F=0*. If, at the extreme, no heterozygotes are observed, then *F=1*. If there is an excess of heterozygotes, F will be less than zero. When *F* is averaged across a large number of independent loci for a population, *F* will typically fall between zero and one. If two populations that differ in allelic profiles are combined (pooled), then Hardy-Weinberg equilibrium is not a sensible null expectation. Divergence of the two allele frequency profiles will manifest as non-random mating and a departure from Hardy-Weinberg equilibrium (the “Wahlund Effect”, Wahlund, 1928). Wright’s *F* applied to the pooled populations is thus informative when examining population subdivision, because the resultant deficit in heterozygotes is an indication of genetic structure. *F* applied in this way incorporates two components -- one arising from departure from Hardy-Weinberg expectation within each population and the second arising from departure from Hardy-Weinberg expectation because of structure between the two populations or sub-populations. For this reason, Wright’s pairwise *F*_*ST*_ is most commonly used as a measure of genetic distance between two populations or sub-populations (see Nei, 1977; Nei & Kumar, 2000:236). *F*_*ST*_ *is* a measure of the heterozygosity attributable to differences in allelic frequency profiles between populations or sub-populations, having partitioned out the contributions from departure from Hardy-Weinberg expectation within populations or sub-populations. Although not a genetic distance in the strict sense, *F*_*ST*_ can be viewed a non-metric distance that varies in value between 0 and 1, such as in studies of isolation by distance. Several methods to estimate *F*_*ST*_ have been developed and some are complex (Excoffier, 2001; Weir & Cockerham, 1984), but there are many software packages available to estimate *F*_*ST*_ for SNP genotype data (e.g. R package hierfstat, Goudet, 2005).

Finally, SNP genotype data at the population level can be converted to binary data by counting the shared alleles between two populations *A* and *B* at a locus

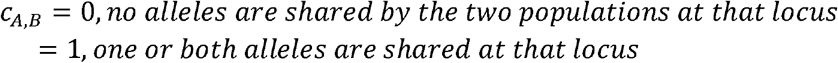

whereby distance measures devised for binary data can be applied. These distances are, in effect, considering only fixed allelic differences between the two populations.

### Visualization of SNP data

There are several methods for visualizing a distance matrix. Perhaps the simplest is to represent the distance matrix as a heat map where colour is used to signify the magnitude of distance between pairs of individuals (Figure 3). Note the white diagonal which indicates zero differences between individuals and themselves, and that the heatmap is redundant in the sense that the lower and upper triangles are identical. A high degree of population structuring is usually immediately evident. For example, individuals of *Emydura* from the north basin of the Murray-Darling catchment are very similar to each other genetically (light blue blocks of Figure 3), but very different from populations of *Emydura* from northern Australia and New Guinea (dark blue blocks of Figure 3). Heatmaps are a very quick way of gaining a visual indication of the level of structuring across the dataset.

**Figure 3.**
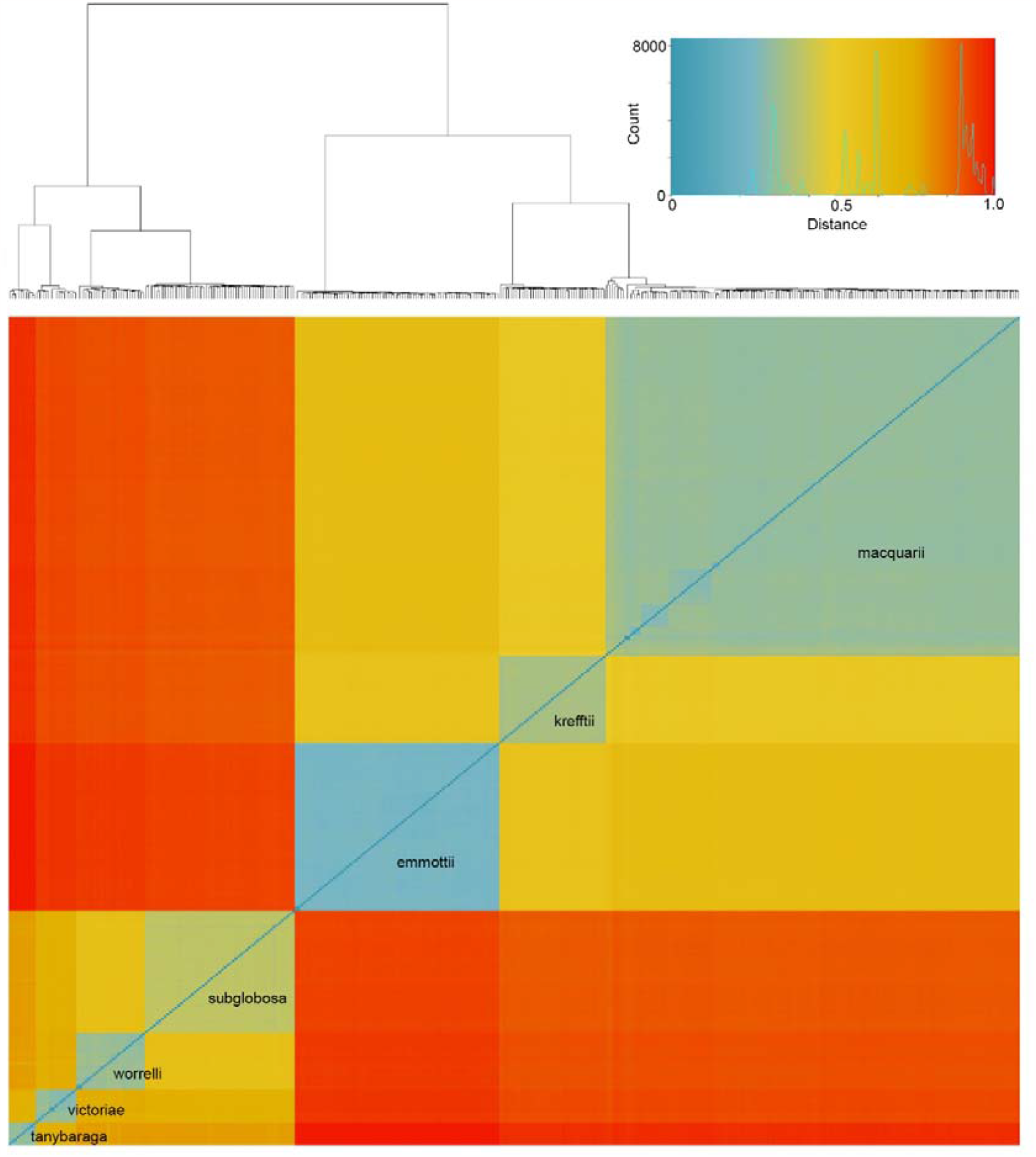
A heatmap used to graphically represent Standard Euclidean Distances calculated pairwise for individual in the freshwater turtle SNP dataset, labelled by sampling locality. Individuals are ranked to highlight clusters. Names of individuals are omitted for clarity. The legend gives the colour scale used to depict the distances with blues for small distances (hence the blue diagonal) and red for the maximum distance of one. The heatmap is accompanied by a neighbour-joining tree. The existence of strong structure is immediately evident. The blocks of low distances aligned along the diagonal represent the putative taxa.

The heatmap is accompanied by a neighbour-joining tree (Saitou & Nei, 1987) which is another common way of graphically representing genetic similarity among entities (Figure 3), not to be confused with their phylogenetic application. An alternative clustering algorithm is UGPMA (unweighted pair group method with arithmetic mean, Sokal & Michener, 1958). Note that to be faithfully represented in tree form, that is, without distortion, the distance measure must be both metric and satisfy the four-point condition (for any four individuals, a branching structure linking them has an internal branch ≥ 0) (Buneman, 1974). This is rarely the case, as real data typically contains conflicting signals as arising, for example, from homoplasy.

By far the most common approach to visualizing genetic distances between entities is Principal Components Analysis (Hotelling, 1933; Jolliffe, 2002; Pearson, 1901) or PCA. PCA is not strictly a distance analysis, but understanding this technique is fundamental to understanding the related Principal Coordinates Analysis (PCoA) which is introduced later. There are at least four computational approaches to calculating a PCA, differing in computational efficiency and tractability (Wu et al., 1997), but essentially PCA takes a data matrix (SNP genotypes or SilicoDArT), represents genetic distance between the entities (individuals or samples or specimens) in a space defined by the *L* loci, centres and realigns that space by linear transformation (rotation) to yield new orthogonal axes ordered on the contribution of variance (represented by their eigenvalues) in their direction (defined by their eigenvector). This process maintains the relative positions of the entities. Because the resultant axes are ordered on the amount of information they contain, the first few axes, preferably the first two or three, tend to contain information on any structure in the data (signal) and later axes tend to contain only noise (Gauch, 1982). This is a powerful visual technique for examining structure in the SNP dataset. An example of a PCA is presented in Figure 4. Important variations include the combination of PCA with discriminant analysis (DAPC, Jombart et al., 2010), adjustment for confounding factors (AC-PCA, Lin et al., 2016) and accommodating sample allelic dropout from low read depth sequence tags (PCAngsd, Meisner & Albrechtsen, 2018). The potential for sNMF (Sparse Non-negative Matrix Factorization), a PCA-related technique useful for disentangling signal from different dyes in microscopy (Rabinovich et al., 2003), may have application in population genetics, particularly for identifying cases of admixture (Frichot et al., 2013). For a broader discussion on variations of PCA not restricted to SNP data, refer to Abegaz et al. (2019). Specific R software for undertaking classical PCA on SNP data and displaying the results graphically includes adegenet (glPCA, Jombart, 2008) and the implementation of adegenet::glPCA in dartR (Gruber et al., 2018; Mijangos et al., 2022), though there is a wide range of packages for undertaking and displaying this very popular analysis.

**Figure 4.**
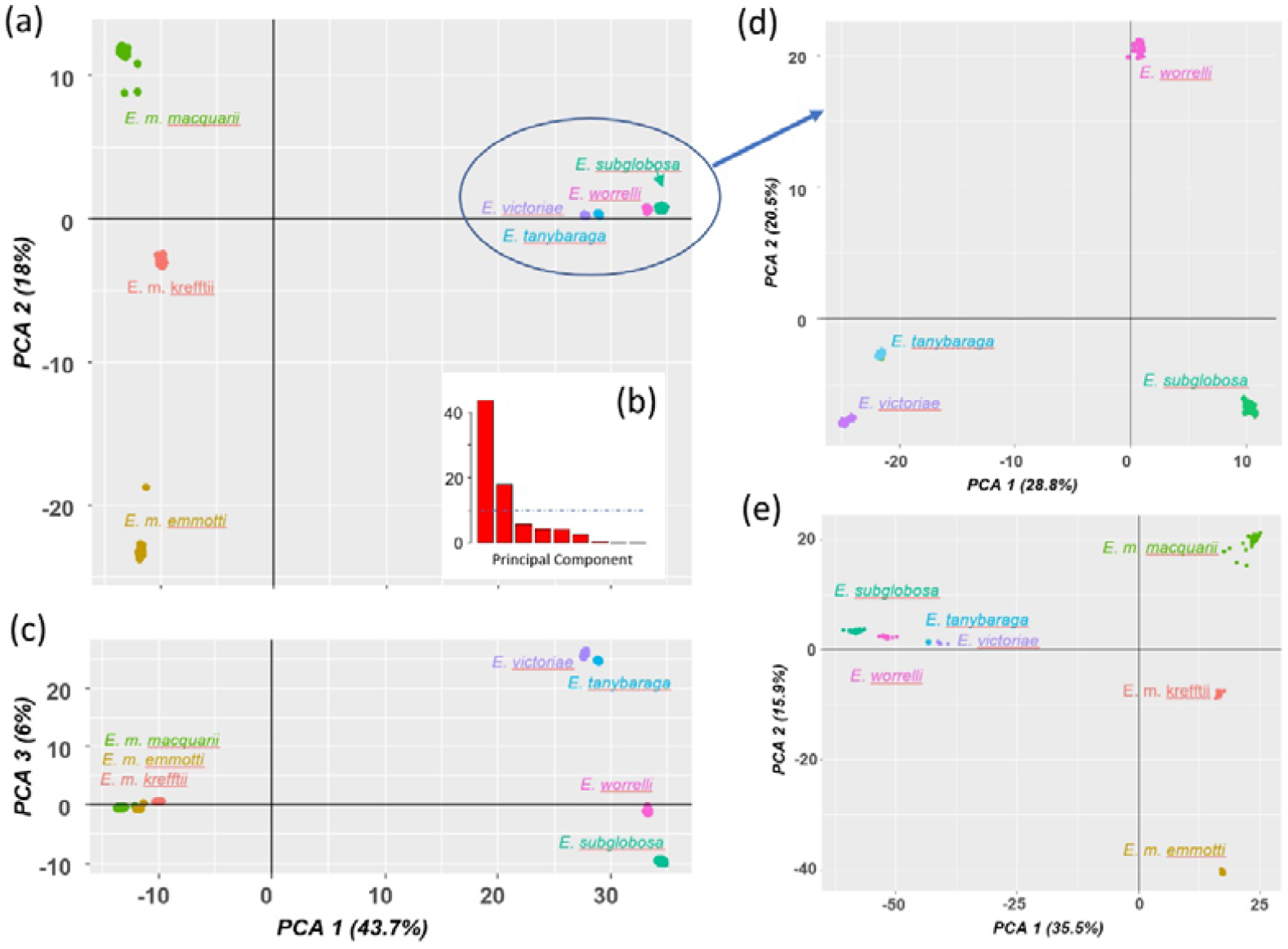
Principal Components Analysis (PCA) as applied to genotype datasets for the Australasian species of *Emydura* (Chelonia: Chelidae). Plot of individuals in a two-dimensional space (**a**) defined by Principal Component 1 (horizontal axis) and Principal Component 2 (vertical axis). Together they explain 61.7% of the total variance among individuals. A scree plot (**b**) shows the contributions to variance by the first nine principal components (those that exceed the Kaiser-Guttman threshold, Guttman, 1954) of which only two each explain more than 10% of variation. Proximity of *Emydura subglobosa/worrelli* to *E. victoriae/tanybaraga* in 2D obscures their distinction, evident when Principal Component 3 is considered (**c**). The variation explained by the first two principal components can be largely set aside by considering structure within the cluster *Emydura subglobosa/worrelli*/*victoriae/tanybaraga* only (**d**), effectively allowing consideration of deeper dimensions. In (e), a PCA using SilicoDArT markers extracts essentially the same structure as the SNP genotype data (albeit flipped horizonally).

Had the data been drawn from a panmictic population (arguably the null proposition), each of the original variables would, on average, be expected to capture the same quantity of variance, and the ordination would fail (the first two axes would each represent only a small and random percentage of the total variance). When there is structure, the percentage of variation explained by the top principal components will dominate the ordination. However, the value of the percentage variation explained cannot be compared across studies as a measure of the strength of the result as if it is some sort of absolute; the percentage variance explained by a principal component needs to be taken in the context of the average percent variation explained by all components (Guttman, 1954). Variances explained by the top 2 or 3 principal components in SNP analyses are typically quite low, but this does not necessarily indicate a poor outcome, as these values need to be considered in the context of the number of axes in the overall ordination.

Note also that a PCA plot can be misleading if one chooses, for convenience, only two or three dimensions in which to visualize the solution. For example, separation in a 2-D plot can be accepted as real but proximity cannot because further separation can occur in deeper dimensions each coupled with a substantial explained variance (Figure 4c). A widely used approach to determining the number of dimensions for the final solution is to examine a scree plot (Cattell, 1966), where the eigenvalues (proportional to variance explained) associated with each of the new ordered dimensions are plotted (Figure 4b). It is usual to apply the Kaiser-Guttman criterion (Guttman, 1954) whereby only those dimensions with more than the average eigenvalue are considered, or to apply a related but less conservative approach to take into account sampling variability (Jolliffe, 1972). Even so, this may result in many informative dimensions. One must decide how much information to discard (say, for example, keeping only those components that explain more than 10% of total variation) or adopt, as a threshold, a visually evident sudden drop in the percentage variation explained, commonly referred to as an elbow. The Kaiser-Guttman criterion is used by dartR to determine how many axes to report in the accompanying scree plot.

More formal techniques (Cangelosi & Goriely, 2007; Jackson, 1993; Peres-Neto et al., 2005) include the broken-stick approach of Macarthur (1957), which provides a good combination of simplicity of calculation and evaluation of suitable dimensionality (Jackson, 1993). The broken-stick model retains components that explain more variance than would be expected by randomly dividing the variance into equal parts. Another related approach is to observe that the eigenvalues of lower “noise” dimensions are likely to decline geometrically, a trend that can be linearized by a log transformation. Informative dimensions are those for which variance explained greatly exceeds that predicted by this linear trend line (more strictly, exert disproportionate leverage on the regression). A more recent approach assesses the statistical significance of the variation explained by each Principal Component (Patterson et al., 2006). Under specified assumptions, the sampling distribution of the ordered eigenvalues, under the null hypothesis of no structure in the data, follows Tracy-Widom distribution (Tracy & Widom, 1993). Thus, it is possible to assign a probability to an observed eigenvalue and retain for consideration only those principal components that are statistically significant (Patterson et al., 2006). The technique is sufficiently robust to violations of its underlying assumptions (e.g. normality) to be applicable to large genetic biallelic data arrays. Application of these more formal techniques require custom R scripts to be prepared by the analyst. Whatever method is used, delineating the informative axes from those representing noise is a personal choice which guides the quantity of signal (as opposed to noise) that is to be discarded in selecting a final visual representation.

### Generalization of PCA

Principal Co-ordinates Analysis (PCoA, Gower, 1966) is an alternative visualization technique that represents a distance matrix in a Euclidean space defined by an ordered set of orthogonal axes, as does PCA. Important variations include adjustment for confounding factors (AC-PCoA, Wang et al., 2022) and application of iterative procedures to best match measured distances against distances in the visual representation for a specified number of dimensions (Belbin, 1991; Kruskal, 1964; Shepard, 1962; Torgerson, 1952). R Software for generating the input distance matrix, then undertaking and displaying graphically a PCoA is provided by ape (function pcoa, Paradis & Schliep, 2019) and the implementation of ape::pcoa in dartR (Gruber et al., 2018; Mijangos et al., 2022).

The primary difference between PCA and PCoA is that PCA works with the covariance (or correlation) matrix derived from the original data whereas PCoA works with any distance matrix. Because the mathematics of PCA moves forward from the covariance (or correlation) matrix, the insight attributed to John Gower (1966) was to substitute any distance matrix for the covariance (or correlation) matrix at this point in the analysis, following a simple transformation. This yields an ordered representation of those distances, metric or otherwise, in multivariable space akin to PCA, greatly expanding the range of application of ordination. In PCA, the *N* individuals are represented in a space of *L* dimensions defined by the loci whereas in PCoA, the individuals are represented in an *N-1* space with coordinate axes based solely on their pairwise distances.

Choosing the number of dimensions to display in visualizing a PCoA is similar to PCA. Missing values are less disruptive for PCoA than classical PCA because they are accommodated in pairwise fashion rather than globally, but they nevertheless require consideration. Missing values result in variation in the precision of estimates of allele frequencies across loci and can break the Euclidean properties of a sample distance matrix even when the chosen metric is Euclidean in theory. Other considerations arise in PCoA from using distance matrices that are non-Euclidean. If a Euclidean distance matrix is used in PCoA then, in the absence of missing values, the distances in the input matrix are represented faithfully in the full ordinated space, that is, without distortion (Figure 5c). The same cannot be said of metric distances in general, as the metric properties ensure that the individuals can be represented in an ordinated space, but not in a rigid linear space (Figure 5a, after Gower, 1982) – some curvature of the space may be required to satisfy the triangle inequality (Figure 5b), which will potentially compromise ability to faithfully represent the distances between entities in a space defined by Cartesian coordinates.

**Figure 5.**
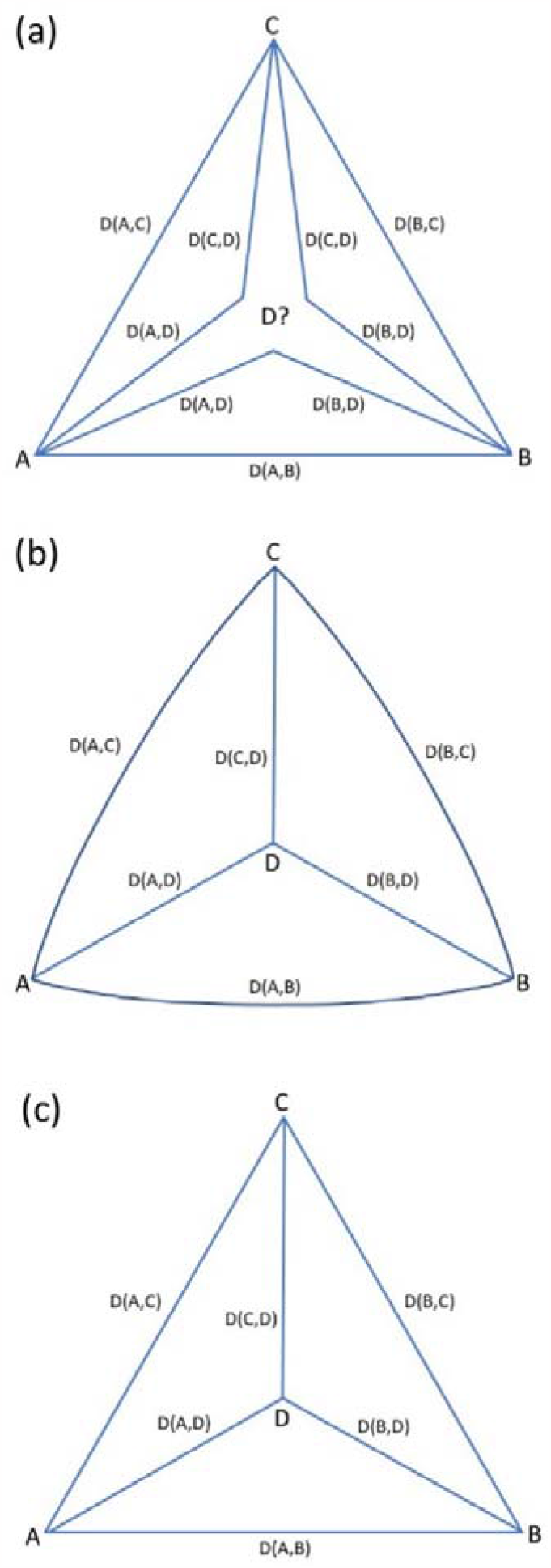
Metricity is not sufficient to represent distances in a rigid space defined by Cartesian coordinates; the distances must be Euclidean. **(a) –** distances between four individuals that satisfy the metric properties can nevertheless not be represented in a linear space defined by Cartesian coordinates because the position of individual D is not defined by the interindividual distances (after Gower, 1982). **(b)** – this distortion can be resolved by allowing non-linear links (geodesics) to represent distances between individuals as, in the case of four points shown, the edges of an irregular Reuleaux tetrahedron in three dimensions. **(c) –** in contrast, Euclidean distances between four individuals can always be represented by linear segments without distortion, as edges of an irregular conventional tetrahedron in three dimensions.

Distortion arising from using non-Euclidean distances manifests as displacement of the points in the visualization, so that the distances among them no longer fundamentally represent the input values; some eigenvalues will be negative (representing imaginary eigenvectors) (Gower & Legendre, 1986). However, the level of distortion is likely to be of concern only if the absolute magnitude of the largest negative eigenvalue is greater than that of any of the dimensions chosen for the reduced representation (Cailliez & Pages, 1976). A few small negative eigenvalues do not detract much from the visual representation if only a few of the highest dimensions are retained in the final solution (Sibson, 1979).

Negative eigenvalues complicate calculation of the variance contributions. In particular, one can no longer calculate the percentage variation explained by a PCoA axis by expressing the value of its eigenvalue over the total sum of the eigenvalues. A correction is necessary (Legendre & Legendre, 2012:506).

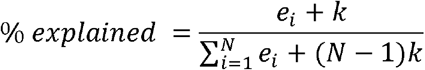

where *e*_*i*_ is the eigenvalue for PCoA axis *i, N* is the number of entities, and *k* is the absolute value of the largest negative eigenvalue.

If negative eigenvalues are considered problematic for the reduced space representation, a transformation can render them all positive and the distance matrix Euclidean. Common transformations put to this purpose are the square root (Legendre & Legendre, 2012:501), Cailliez transformation (Cailliez, 1983; Gower & Legendre, 1986) and the Lingoes transformation (Gower & Legendre, 1986; Lingoes, 1971).

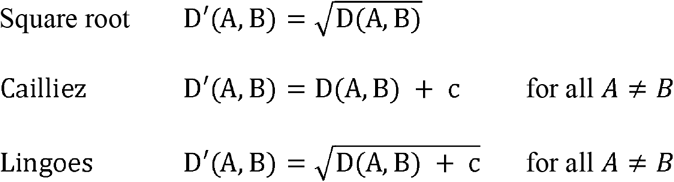

The value of *c* is chosen to be the smallest value required to convert the most extreme negative eigenvalue to zero.

Finally, the outcome of a PCoA with an input matrix comprised of Standard Euclidean Distances calculated from individual genotypes is identical to the outcome of a PCA (Cox & Cox, 2001:43-44). The interchangeability of the two, PCA and PCoA, in this restricted case leads to confusion on the distinction between the two analyses.

Notwithstanding that the final arrangement of entities in the ordinated space is the same for PCA and PCoA using Euclidean Distance on the same data, the percentage variation explained by each axis will differ between the two approaches. This is because PCA calculates the percentage variation explained by an axis as the percentage its eigenvalue represents of the total of all eigenvalues. The eigenvalues are calculated from the covariances generated from the input data matrix. PCoA in contrast will calculate the percentage variation explained by an axis as the sums of squares of the distance between the entities in the direction of the axis expressed as a percentage of the total sums of squares of the distances in the input distance matrix. The two will differ even when PCA and PCoA using Euclidean Distance are applied to the same data. Hence, it is not possible to simply compare the performance of a PCA and a PCoA based on examination of the values of percentage variation explained.

### Missing values

SNP genotype datasets typically have substantial numbers of missing values. Missing data can arise because the read depth is insufficient to detect SNP-containing sequence tags consistently, or because mutations at one or both of the restriction enzyme recognition sites in some individuals result in allelic drop-out (null alleles). Because those that arise from mutation are inherited, they are subject to genetic drift (and potentially local selection) differentially within each sampled population, and so are not randomly distributed across the entire dataset. Indeed, in some populations the mutation(s) leading to missing values may become fixed. Filtering on call rate with a threshold applied globally will potentially admit high frequencies of missing data at particular loci for single populations, which has consequences as outlined below. In SilicoDArT data, the null alleles are themselves the data, scored as presence or absence of the amplified tag. In this context, the marker data extraction pipeline has to apply adequate quality control of read depth, because hemizygous individuals are scored as present (1).

Missing data are problematic. Techniques like classical PCA that access the raw data matrix cannot accommodate missing data. PCoA, which accesses a distance matrix, is less affected by missing values in the raw data (Rohlf, 1972) because they are eliminated in pair-wise fashion during construction of the distance matrix, not globally. PCoA is nevertheless affected by missing values in ways that are not immediately transparent – by potential breakage of metric or Euclidean properties, and by disregarding variable precision of estimates of population allele frequencies across loci. The trade-off is one of balancing data loss with bringing distortion of the visual representation within acceptable levels.

Because classical PCA will not accept missing values, when a locus is not scored for a particular individual, either the data for the entire locus must be deleted or the data for the entire individual must be deleted. This is clearly very wasteful of information, and unacceptable loss when working with SNP genotype data. The degree of data loss can be controlled to an extent by pre-filtering on call rate by locus (say requiring a call rate > 95%) or by individual (say requiring a call rate > 80%), noting that a missing value rate of <10% causes only modest overall distortion using implicit mean-imputed missing data (Figure 6a). Numerous ways for handling missing data in PCA have been suggested (Dray & Josse, 2015), but the most common method is to replace a missing value at a locus with the mean of the allele frequencies for that locus calculated across the individuals for which it is non-missing (mean-imputed missing data). Mean-imputation of missing data can lead to the individuals with a high proportion of missing data being drawn out of their natural grouping and toward the origin, leading to potential misinterpretation (Yi & Latch, 2021). For example, if individuals in the PCA aggregate into natural clusters, perhaps representing geographic isolates, and these clusters are on either side of the origin, then an individual with a high frequency of missing values will be drawn out of its cluster when the missing values are replaced by global average allele frequencies (Figure 6). Its location intermediate to the two clusters might be falsely interpreted as a case of admixture. Individuals with missing data corrected by mean-imputation will also distort confidence envelopes as applied to clusters with consequences for interpretation, particularly in studies of population assignment (Figure 7).

**Figure 6.**
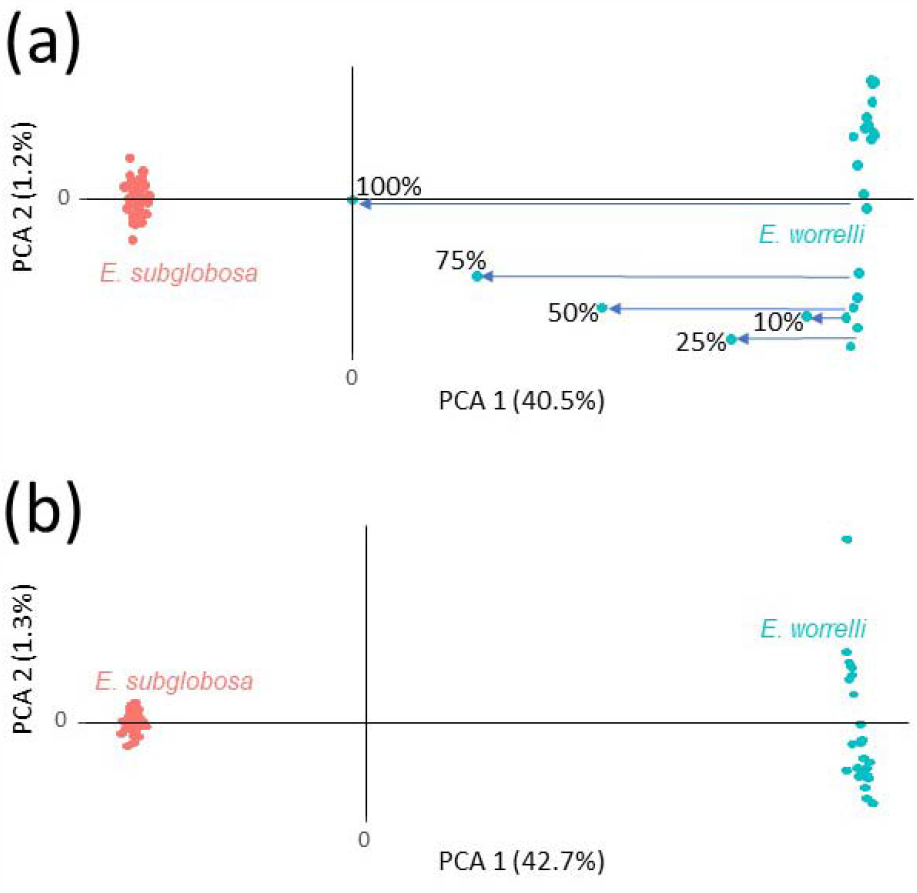
Principal Components Analysis (PCA) applied to two populations where five individuals in one population (*Emydura worrelli*) have had percentages of SNP loci ranging from 10% to 100% set to missing. **(a)** – the impact of this on PCA where missing data are filled with the average global allele frequencies (mean-imputed missing data) is clear, and subject to misinterpretation as hybridization or various levels of introgression. **(b)** – the issue can be resolved by local imputation, in this case by nearest-neighbour imputation.

**Figure 7.**
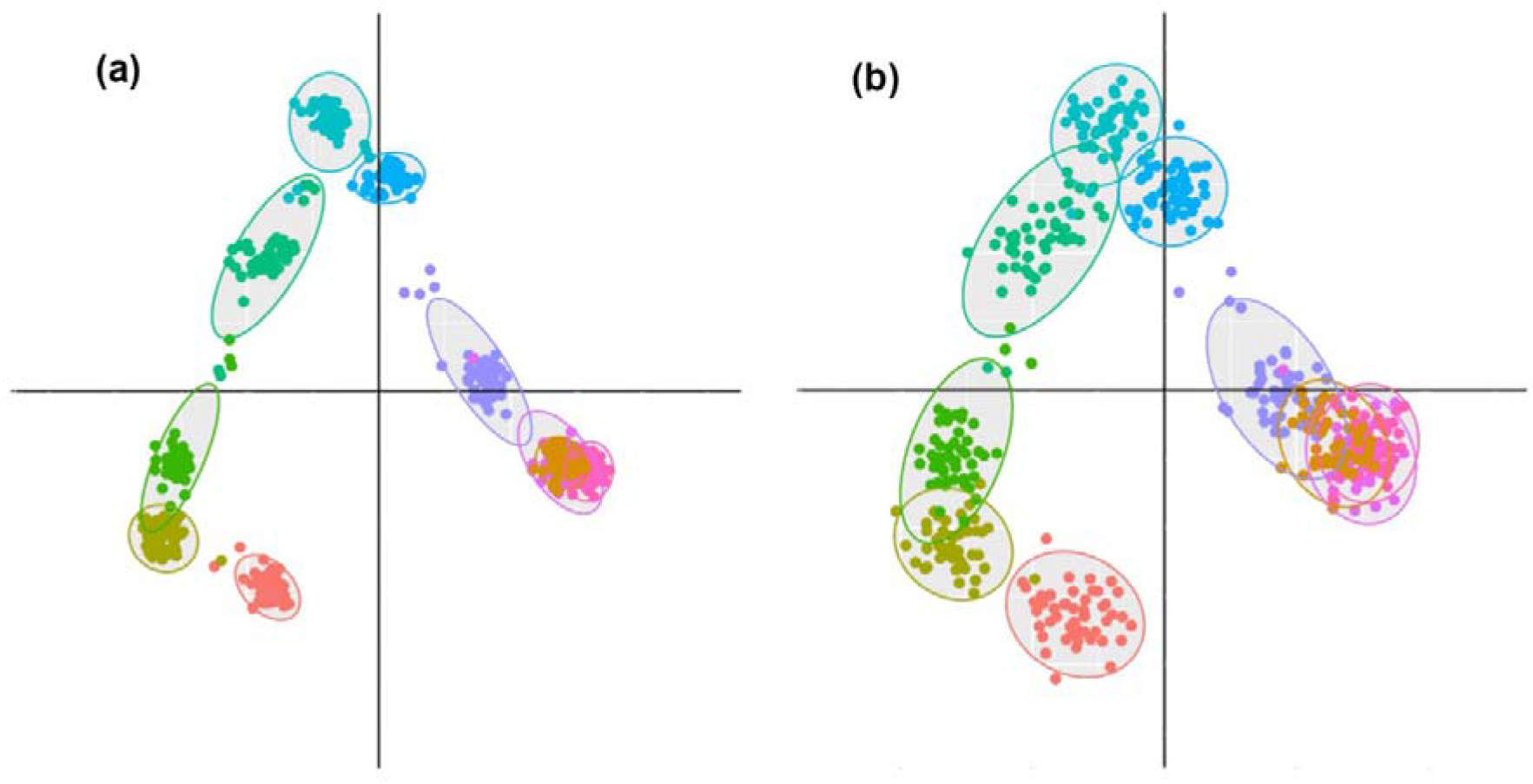
A PCA plot of simulated data showing aggregations and their associated 95% confidence limits for (a) data with no missing values and (b) data with 50% missing values. Note the inflationary effect missing values have on the spread of the points in each aggregation, manifested as inflation of the 95% confidence limits, an artifact. This is of particular relevance to studies of population assignment.

Several better ways than global mean-imputation for handling missing values in SNP are appropriate:

1. Replace missing values for an individual with the mean observed allele frequency for the population from which the individual was drawn. In this way, the individual is displaced toward the centroid of the population from which it was drawn, not the origin of the PCA.
2. Replace missing values for an individual with the expected value for the population to which the individual belongs, based on the assumption of Hardy-Weinberg equilibrium. The individual will again be displaced toward the centroid of the population from which it was drawn, not the origin of the PCA.
3. Replace missing values for an individual by the value at the same locus from its nearest neighbour (the individual closest to the focal individual based on Standard Euclidean genetic distance) (Beretta & Santaniello, 2016) (Figure 6). The focal individual will be drawn toward its nearest neighbour, typically an individual within the same population. This method has the advantage of filling missing values even where all individuals collected from a single location are missing for a given locus.
4. Generate a Euclidean distance matrix with pairwise deletion of missing values (rather than global deletion required of PCA) and apply PCoA to generate the ordination. This relies on the broad equivalence of PCA and PCoA using Euclidean distance (Cox & Cox, 2001:43-44).

For SilicoDArT, because missing values are typically associated with hemizygous individuals, imputation can be achieved by recoding the missing values as 1.

### Linkage

Distance analyses usually assume that each locus contributes independent information to the overall distance value (i.e. they segregate independently). With the large genotype arrays typical of SNP datasets, some SNP loci are likely to be linked (Waples et al., 2022). In extreme cases, linkage disequilibrium can seriously distort the genetic structure, confounding interpretation. Large blocks of sequence with limited haplotype variation (presumably a result of limited recombination) have been observed in humans (Daly et al., 2001; Patil et al., 2001) and plants (Battlay et al., 2022) and their impact modelled in simulations (Lotterhos, 2019). If many markers have been sequenced in such a haploblock, they will be tightly linked, and support for any population structure that they represent will be proportionately inflated. This will be evident in a PCA or PCoA as artifactual structure that will potentially defy explanation when working with organisms with little genomic information. An example of such artifactual structure in a PCA plot generated from linked SNP markers is provided by polymorphism in a large inversion in human chromosome 8 (Amos & Ma, 2012; Battlay et al., 2022). SNP markers associated with this inversion generate a coordinated signal which manifests as a three-group pattern, one corresponding to inverted homozygotes, one corresponding to heterozygotes and one corresponding to non-inverted homozygotes (refer also to the simulations of Lotterhos, 2019). Such an explanation was invoked to explain the disaggregation of individuals of the Australian dragon *Pogona vitticeps* into two mirrored clusters that could not be explained by location of capture, sex effects or batch effects (Wild et al., 2022). The signature three-group pattern characteristic of a large polymorphic inversion was evident also in a SNP study of genetic variation across swamps occupied by the Australian Blue Mountains Water Skink *Eulamprus leuraensis* (Figure 8).

It is wise as a data quality step to prune groups of SNPs with high linkage disequilibrium (LD). In extreme cases of linkage arising from SNPs residing on a shared, non-recombining block in the genome (Figure 8) can be resolved by identifying loci that are strongly correlated with the Principal Component that separates out the artifactual groupings and removing those loci from the analysis (Wild et al., 2022) or by SNP pruning (Lotterhos, 2019) albeit with substantial loss of data. The approach of Wild et al. (2022) can also be used to remove batch effects (Lou and Therkildsen, 2022) when they dominate the PCA visualization as an alternative to more intrusive approaches such as removing all private alleles (Lou and Therkildsen, 2022). Batch effects are best managed appropriate quality control at the time of generating the SNPs and SilicoDArT data. It is important also to ensure that SNP and SilicoDArT scores are generated together and the analysis rerun for the combined sequences rather than attempting to combine datasets posthoc.

**Figure 8.**
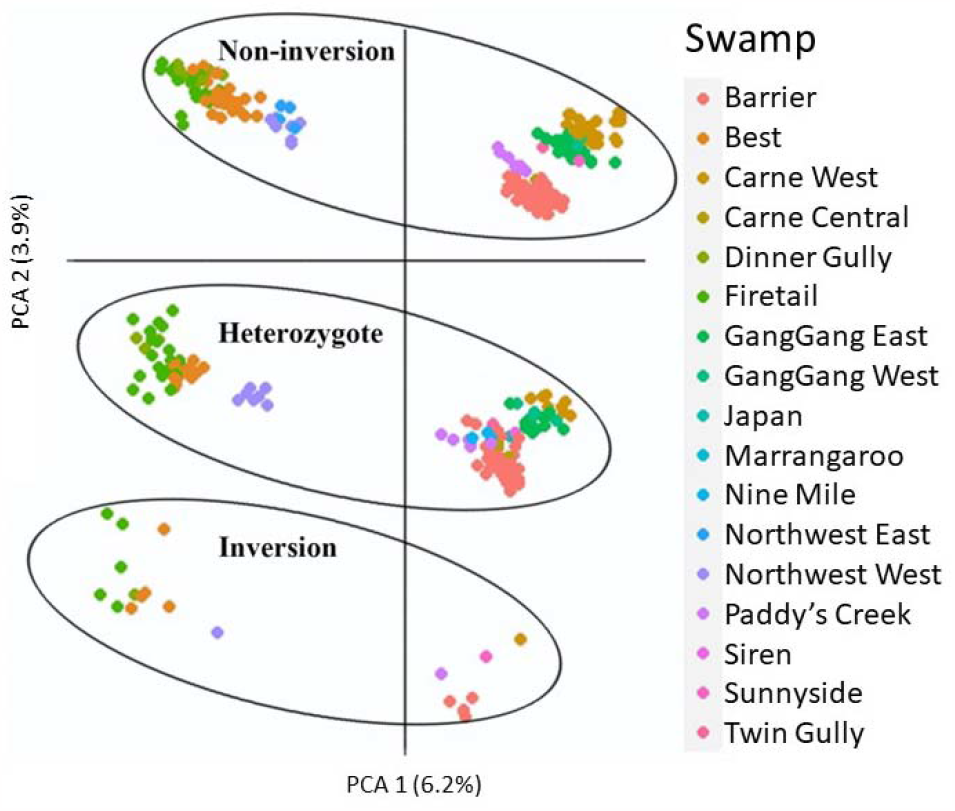
An ordination of SNP genotypes for the Australian Blue Mountains Water Skink *Eulamprus leuraensis* collected from 17 locations with varying levels of isolation. The ordination shows the three-group structure characteristic of a large autosomal inversion (Amos & Ma, 2012). For sake of illustration, we have assumed that the inversion is the least frequent of the two polymorphic states. This interpretation remains speculative until sufficient genomic information is available for this species to demonstrate the existence of the inversion and to associate the SNP loci strongly correlated with PCA 2 to that inversion.

### Relatedness among individuals

Representing distance between populations on the basis of samples typically assumes that the individuals are sampled at random and their genotypes are independent. Systematic errors can arise if some sampled individuals are more closely related than are individuals selected at random from their population (Östergren et al., 2020); these individuals can be expected to separate out from the main body of individuals in a PCA or PCoA (Figure 9a). This can occur if, for example, individuals are selected from a single school of fish that may be more closely related than individuals chosen at random. The effect is most pronounced if parents and their offspring (siblings) are among the sampled individuals (Figure 9a). Care should also be exercised when interpreting clinal structure if the unit of dispersal comprises groups of related individuals (Fix, 1997). These closely related individuals should be identified and all but one removed from the analysis, with a close eye to the consequences of overzealous removal of related individuals (Waples and Anderson, 2017).

**Figure 9.**
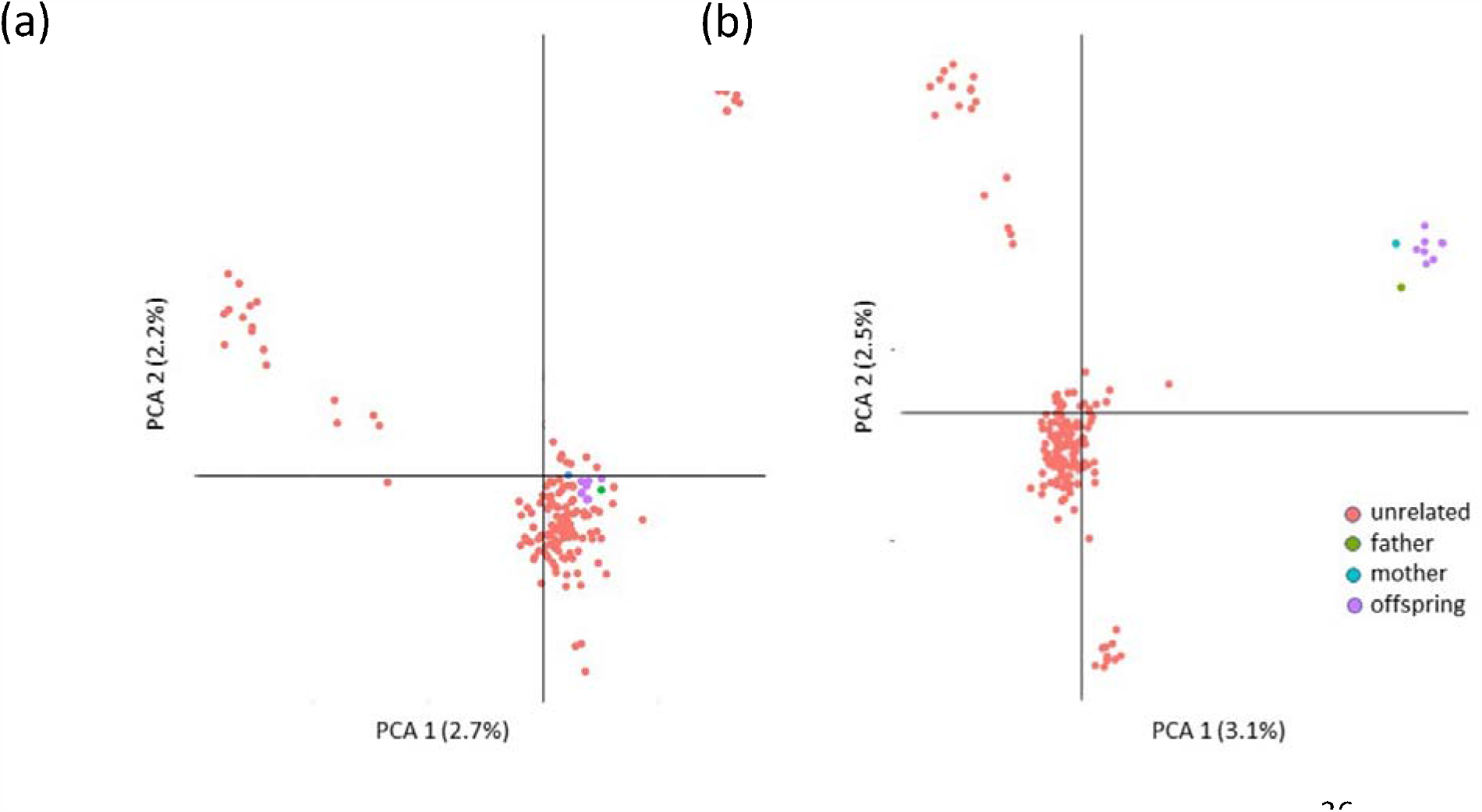
A series of sub-populations of *Emydura macquarii* from the northern basin of the Murray-Darling drainage (a) in which one subpopulation (central) has two individuals identified to be manipulated artificially into a parent-offspring relationship and eight individuals into a parent-offspring relationship; (b) The ordination after the addition of the modified individuals. If such closely related individuals go undetected, and are retained, the spatio-temporal distance relationships between populations can be subject to misinterpretation.

## Discussion

Species and populations usually do not constitute a single panmictic unit where individuals breed at random over their entire range. Population subdivisions typically exist, which may be hierarchically arranged if they reflect patterns of ancestry and descent, or not if they are each on independent random walks, perhaps periodically reset by episodic gene flow (Georges et al., 2018). It follows that patterns evident in distances between entities, be they individuals, aggregations of individuals, populations or taxa, will often be highly structured. The challenge for SNP analyses arises from dealing with an incredible volume of data, certainly in the thousands of markers but often in the hundreds of thousands or even millions of markers, each with its own contribution to overall variation. Summarising the data as a distance matrix and visualizing the important structure using ordination is a first step in examining structure and hypothesising on its potential causes.

Many distance measures are available for different types of genetic data (Kosman & Leonard, 2005; Libiger et al., 2009). Fortunately, characteristics of SNP data and considering contemporary rather than historical processes governing variation among populations, dramatically reduce the options from which to select an appropriate distance. SNP markers are biallelic in practice, so distinct distance measures in a multiallelic context can become the same in a biallelic context. Additionally, symmetry considerations in the arbitrary choice of reference and alternate allele eliminate use of distance measures that treat scores of homozygous reference (0) as fundamentally different from scores of homozygous alternate (2). Because genetic distances are likely to be interpreted in the context of everyday notions of a distance measure, even if subconsciously, distances that satisfy the basic metric properties are preferable over those that do not. The decision on which distance measure to use then comes down to one of context and whether or not underlying evolutionary models of mutation and drift are considered important.

Genetic structure within and among populations is commonly explored using ordination which is inherently reliant on the concept of genetic distance. Strictly, distances with Euclidean properties are required to faithfully represent those distances in a space defined by Cartesian coordinates using PCA and PCoA. Distances with metric properties yield adequate representations in most cases (e.g. Jaccard D, Manhattan D, Chord D). Non-metric distances should in principle be avoided when the goal is to undertake a distance analysis and ordination, notwithstanding their value in a broader context (e.g. Fst). However, methods for detecting distortion of a visual representation arising from gross departures in Euclidean and non-metric properties (negative Eigenvalues) are available as are means for redressing the deficiency. In practice, the distortion arising from selecting only the top two or three axes for the visualization generates more distortion than does a relatively few negative eigenvalues. One can thus be somewhat relaxed about departures from Euclidean or metric properties of a distance if there are compelling reasons for choosing a non-metric distance measure in the context of the study, one’s philosophical position on the need for an underlying evolutionary model and questions to be explored.

PCA has recently come under criticism (Elhaik, 2022). The deficiencies of PCA identified by Elhaik are real, but we argue that they derive from over-interpretation of PCA to address questions that are beyond its remit, and because of misunderstanding of what ordination delivers as its ultimate solution. PCA is essentially an exploratory tool that draws out patterns from multivariable data by aggregating correlated influences into a relatively few dimensions. PCA is about extracting patterns. It has little to contribute to understanding of the processes that delivered those patterns however tempting it might be to use the visualizations to make inferences about underlying events and processes (McVean, 2009).

The patterns thus are suggestive but not demonstrative of underlying causes. Interpretations of the causes of the patterns need to be pursued by more definitive subsequent analysis to distinguish between competing explanations for the observed patterns (Novembre and Stephens, 2008). For example, individuals that fall in an intermediate position between two well defined groupings in a bivariate plot of PCA1 and 2 may be an indication of admixture. Certainly, that is the position in the ordination plot that admixed individuals will fall, but the reverse inference is not true. The intermediary position may not be sustained on examination of deeper dimensions or may have arisen from other causes. Hybridization and introgression are complex processes that cannot be simply inferred from an examination of the patterns evident in a PCA (Gompert and Buerkle, 2016). Definitive demonstration of admixture requires additional analyses (refer Figure 4 of Georges et al., 2018), such as the frequency of heterozygotes at loci fixed and different for the two major groupings, or use of statistical packages such as NewHybrids (Anderson & Thompson, 2002). Similarly, PCA may be undertaken as a prelude to genome-wide association studies (GWAS) to identify genes under selection (Xu et al., 2022). Serious difficulties arise in terms of defensible and reproducible conclusions when interpretation is restricted to PCA without targeted and appropriate analyses to follow the indications from the PCA. One cannot reliably use two or three-dimensional representations in PCA alone to decide ancestry or demographic history, to reliably identify and exclude outliers, to infer kinship, to identify ancestral clines in the data, to detect genomic signatures of natural selection, or to identify cases of convergent evolution (Elhaik, 2022 and references cited therein). Misleading artifacts can emerge if one attempts to interpolate a surface between observed sample points in the ordinated space, artifacts such as sinusoidal trends or ‘waves’ that can lead to misinterpretation of migration patterns (Novembre and Stephens, 2008). PCA and PCoA can provide only suggestive indications useful for formulating hypotheses for further rigorous examination. Therein lies the value of ordination.

A second criticism of PCA is that the visual representation of distances between individuals is sensitive to sample size (Elhaik, 2022). Small sample sizes will have a greater associated error in their representation of the population from which they are drawn, and this will be reflected in the outcome of the ordination. But the sample size dependency identified by Elhaik arises because decreasing the sample size of one structural grouping (population, aggregation of populations, taxon) will reduce its contribution to overall variance and reorient the individuals in the space defined by the informative dimensions of the ordination. This leads to potentially dramatic changes in their projection on to the top two dimensions. Thus sample size effect can have major consequences for interpretation if the analyst’s focus is restricted to two or three-dimensional PCA plots.

The apparent lack of reproducibility identified by Elhaik (2022) arises from restricting attention to only the top two (or three) of informative dimensions. Misinerpretation arising from this effect can be overcome by appropriate examination of the ordination, including all substantial informative dimensions.

Arguably, missing values, and how they are managed, is the issue with the most significant consequences for genetic distances and their visualisation. Classical PCA requires a complete input dataset, and adequate filtering missing values requires removal of loci or individuals from the analysis to create the input matrix free of missing values. For SNP data, which typically have high frequencies of missing scores, the consequent data loss is unacceptable. To overcome unacceptable loss of data, the missing values are typically infilled using the global average for the locus concerned. We have shown that this can result in misleading displacement of individuals with high rates of missing data away from their natural groupings and toward the global centroid, and inflation of confidence envelopes for aggregations of individuals with the risk of misinterpretation, particularly in analyses such as population assignment. We have reviewed the approaches to deal with missing values that avoid these distortions, the most effective of which appears to be replacement of a missing value with that of the nearest neighbour (Beretta & Santaniello, 2016). Even replacement with a random value is preferable to replacement with the global average allele frequency (the most common default approach) because random values simply add noise to the data which is driven to lower dimensions in any ordination.

Other concerns are linkage, which is difficult to detect and filter when sample sizes from collection localities are small and cannot be pooled because genetic structure among collection localities contributes artificially to estimates of linkage disequilibrium. In gross cases of linkage, such as those arising from a polymorphic chromosomal inversion (Figure 8), SNPs associated with the inversion can be identified and removed by screening out loci that load high with the PCA axis most affected by the inversion (Wild et al., 2022) or by SNP thinning (Lotterhos, 2019) albeit with substantial loss of data.

In summary, we recommend selection of a distance measure for SNP genotype data that does not give differing outcomes depending on the arbitrary choice of reference and alternate alleles in the case of SNP data, and careful consideration of which state should be considered as zero when applying binary distance measures to SilicoDArT data. As a broad recommendation on choice of a distance measure, Standard Euclidean Distance is the default choice of a genetic distance for SNP and SilocoDArT data. It is free of assumptions about the processes that generate the variation and, in the biallelic case, does not suffer the unacceptable distortions that arise in multiallelic data when two populations are polymorphic at a site, but share no alleles (Nei & Kumar, 2000:246). Standard Euclidean Distance can be faithfully represented in ordinated Cartesian space. Other metric distances may be chosen in order to down weight joint absences in the case of SilicoDArT data (e.g. Jaccard Distance) or because of the importance of a particular model of progressive divergence with time (e.g. Nei’s D, Reynold’s D, Chord D) for SNP genotype data. For example, Chord Distance is better correlated with time since divergence and is a metric distance, so may be a better choice than Fst /(1-Fst) (a non-metric distance) when analysing isolation by distance (Séré et al., 2017). Diagnostic examination of eigenvalues should be undertaken when a non-metric distance has been selected, or if the analysis is to include substantial missing values which breaks metricity. Action should be taken if the sum of negative eigenvalues is substantial to avoid distortion in the final visual representation. We strongly recommend filtering heavily on missing values, then imputing those that remain to create a full matrix prior to undertaking a distance analysis. Failure to do so can substantially and artificially inflate confidence envelopes for populations or aggregations of populations and lead to other distortions that can lead to misinterpretation. Screening for closely related individuals (parent-offspring or sibling relationships) is also important, and the impact of polymorphic haploblocks in the genomes of target species or populations occasionally emerges as a challenge for studies of species with limited genomic information.

## Conflict of Interest

Author AK is the Managing Director of Diversity Arrays Technology Pty Ltd (Canberra) and the originator of the DArTSeq pipelines and associated data formats (including SilicoDArT). The other authors have no conflicts of interest to declare.

## Data Availability

Real datasets and the scripts used to generate simulated datasets used in this paper are available on Dryad (https://doi.org/10.5061/dryad.4b8gthtkn).

## Author Contributions

All authors contributed to the development of ideas presented in this manuscript. AG led the writing, LM and BG undertook the simulations, HP provided input on human studies, MA provided access to the skink data and provided associated context.

## Acknowledgements

We would like to thank Bill Sherwin, Fred Allendorf, Evsey Kosman and anonymous reviewers for comments on an earlier draft of this manuscript, and for assisting with setting the scope and direction of the manuscript. We would like also to thank Springvale Coal Pty Ltd for allowing the publication of genetic data obtained for the Blue Mountains Water Skink Research and Management Program coordinated by RPS Australia East Pty Ltd for the Springvale Extension Project. This work was funded in part by contributions from The Cooperative Research Network for Murray-Darling Basin Futures, Australian Department of Industry, Innovation, Science, Research and Tertiary Education (DIISRTE). Grant Code: MDB-CRN (2012-2014); ARC Linkage Program: Drivers of fine scale genetic spatial structuring in aquatic organisms, Australian Research Council LP140100521 (2014-2017); and the ACT Priority Investment Program: EcoKDDart Future Proofing Biodiversity and Biosecurity with a novel integrative Big Data Analysis Hub, ACT Department of Territories and Local Government, Grant 638238 (2020-2023); via their contributions to the development of dartR.

## Notes

https://doi.org/10.5061/dryad.4b8gthtkn

### Summary of Updates

This revision has been substantially revised in response to comments from reviewers and others re a new submission. An additional author has been added and better treatment of SilicoDArT datasets is included. A link to datasets and R scripts to replicate the analyses have been added.

